# Large Extracellular Vesicles Transfer Higher Levels of Prions and Infect Cell Culture Better than Small Extracellular Vesicles

**DOI:** 10.1101/2023.03.01.530593

**Authors:** Jakub Soukup, Tibor Moško, Sami Kereïche, Karel Holada

**Affiliations:** Institute of Immunology and Microbiology, First Faculty of Medicine, Charles University, 128 00 Prague, Czech Republic; Department of Genetics and Microbiology, Faculty of Science, Charles University, 128 44 Prague, Czech Republic; Institute of Biology and Medical Genetics, First Faculty of Medicine, Charles University, 128 00 Prague, Czech Republic

**Keywords:** Prion, PrP^TSE^, Extracellular Vesicles, Cell Culture, Infection

## Abstract

Prions are responsible for a number of lethal neurodegenerative and transmissible diseases in humans and animals. Extracellular vesicles, especially small exosomes, have been extensively studied in connection with various diseases. In contrast, larger microvesicles are often overlooked. In this work, we compared the ability of large extracellular vesicles (lEVs) and small extracellular vesicles (sEVs) to spread prions in cell culture. We utilized CAD5 cell culture model of prion infection and isolated lEVs by 20,000 × g force and sEVs by 110,000 × g force. The lEV fraction was enriched in β-1 integrin with a vesicle size starting at 100 nm. The fraction of sEVs was depleted of β-1 integrin with a mean size of 79 nm. Both fractions were enriched in prion protein, but the lEVs contained a higher prion-converting activity. In addition, lEV infection led to stronger prion signals in both cell cultures, as detected by cell and western blotting. These results were verified on N2a-PK1 cell culture. Our data suggest the importance of lEVs in the trafficking and spread of prions over extensively studied small EVs.

## Introduction

Prion diseases, such as CJD in humans or scrapie and bovine spongiform encephalopathy in animals, are connected with the accumulation of pathologically misfolded prion protein (PrP^TSE^) in the brains of affected individuals [1]. Normal cellular prion protein (PrP^C^) has various posttranslational modifications, including two potential N-glycosylation sites, which can be recognized by western blotting as three distinct bands [2,3]. PrP^C^ is localized on membranes by a C-terminal GPI anchor [4]. Its structure contains three alpha helixes and a short beta-sheet [5,6]. The misfolding of PrP^C^ into PrP^TSE^ can occur sporadically, with genetic predisposition [1], induced to PrP^C^ by contact with PrP^TSE^ [1]. The beta-sheet structure of PrP^TSE^ is resistant (PrPres) to digestion with proteinase K (PK) [7], which could be used to track infection in organs or cell cultures. Prions are most likely transmitted horizontally between individuals by the oral pathway [8]. However, how exactly prions spread from the gut to the brain remains unknown. One possible theory described by McBride et al. [9] is that prions could transmit via splanchnic and vagus nerves. Other possible routes are through lymphoid tissue [10] and follicular dendritic cells [11]. Intercellular transmission occurs in three possible ways: 1) cell-to-cell contact [12], 2) tunneling nanotubes [13], and 3) EVs [14]. Other mechanisms have been proposed, such as trogocytosis or GPI-painting [15,16].

Extracellular vesicles (EVs) are small cell-derived lipid vesicles with significant physiological and pathological functions [17]. EVs can be divided into three primary groups depending on their size and biogenesis. Exosomes are generated by endosomal sorting complex required for transport (ESCRT)-dependent [18,19] and ESCRT-independent pathways [20]. The ESCRT-dependent pathway involves the ESCRT mechanism, formed by four distinct complexes (ESCRT-0, ESCRT-I, ESCRT-II, and ESCRT-III) of inward budding in early endosomes generating intraluminal vesicles (ILVs) and maturing early endosomes in multivesicular bodies (MVBs) [18,19]. MVBs can be recycled back to the cell membrane, releasing ILVs as exosomes with sizes of 30–100 nm [18,21]. Another pathway leads MVBs to fuse with lysosomes, degrading ILVs [22,23]. Additionally, MVBs can fuse with autophagosomes, creating amphisomes from which ILVs are released [24,25]. As a result, exosomes contain ESCRT-associated proteins such as TSG-101, tetraspanins (CD9, CD63, CD81), or Alix (through the Syntenin-Alix pathway [26,27]). An ESCRT-independent pathway utilizes neutral sphingomyelinase (NSMase), which hydrolyzes sphingomyelin to ceramide and induces the aggregation of microdomains to form ILVs [20].

Microvesicles (MVs) are vesicles 100–1000 nm in size [18] generated by direct budding from the cell membrane. The budding of MVs is regulated by Cdc42 (a Rho family protein) and its downstream effector Ras GTPase activating-like protein 1 [28]. The actin cytoskeleton regulator RhoA is also involved in the dynamics of MV budding [29,30]. ESCRT-I and III mechanisms also play a role in MV biogenesis. Furthermore, ceramide has been reported to be involved in the biogenesis of MVs [31,32]. Cone shape-structured tetraspanins are also involved in forming MVs [33]. Based on their biogenesis, MVs contain adhesion molecules such as β-1 integrin [34,35] or Annexin A1 [36]. The last primary type of extracellular vesicles is apoptotic bodies, which have variable sizes between 500 and 5000 nm and are generated during apoptosis [35]. Apoptotic bodies are enriched with phosphatidylserine on their surface, targeting it for fast phagocytosis by leukocytes [35], which should eliminate them from effective cellular communication [37]. Recently, more subpopulations of EVs like small exomeres or large oncosomes have been distinguished using various isolation techniques [27,36,38,39].

In the cell, PrP^C^ is translocated to the plasma membrane in both nonraft [40] and raft domains [41–43], where it is recycled back by clathrin- or caveolin-1-coated pits to endosomes, from which it can mature into MVBs [40,44–46]. PrP^C^ is prevalent in MVBs [47,48], with endolysosomal complexes serving as one of the conversion sites for PrP^TSE^ [49,50]. Vilette et al. [51] found that exosomal prion infectivity is associated with both ESCRT-dependent and ESCRT-independent release pathways. The intercellular infectivity of PrP^TSE^ is decreased by blocking NSMase-2, while the PrP^TSE^ content in the cell increases [48,52]. PrP^TSE^ present in exosomes [14] is able to infect neuronal cells both in vitro and in vivo [53]. One of the possible physiological roles of PrP^C^ is the mediation of EV transport along neuronal cells, for which PrP^C^ located in EVs hydrophilically interacts with neuronal PrP^C^ located on the plasma membrane [54], which is proposed to be the primary conversion site for PrP^TSE^ [49,55]. From the plasma membrane, PrP^TSE^ is also transported via the endolysosomal pathway [56] to lysosomes [57] and possibly MVBs. As in exosomes, PrP^TSE^ on membrane-derived microvesicles is also released from cells [58].

PrP^TSE^ is present in plasma EVs of symptomatic and asymptomatic mice infected with prions [59,60]. Thus, via blood transfusion, EVs can contribute to recorded horizontal infection with variant CJD [61]. The involvement of EVs in the oral pathway of infection is also possible since EVs can survive the stomach environment [62,63].

In our work, we utilized CAD5 cell culture model of prion infection to compare PrP^TSE^ content and prion infectivity in the sEV fraction and the lEV fraction. According to MISEV 2018 [64], we describe our EV fractions as lEVs pelleted by 20,000× g and enriched in β-1 integrin and sEVs pelleted by 110,000× g and depleted by β-1 integrin. Our findings were similar on N2a-PK1 cell culture model with similar methodology, although the differences were lower. Here, we report that both fractions of EVs are enriched in PrP, but lEVs are able to infect native cells with more efficiency than sEVs.

## Methods

### Materials

Antibodies: From Abcam, we obtained rabbit anti-β-actin (EPR21241), fin. con. 0.25 µg/ml; and mouse anti-Alix (3A9), fin. con. 2 µg/ml. From Azure Biosystems, we obtained goat anti-mouse AzureSpectra 700 or 800 conjugate, fin. con. 0.4 µg/ml; and goat anti-rabbit AzureSpectra IR700 or IR800 conjugate, fin. con. 0.4 µg/ml. From Enzo Life Sciences, we obtained rabbit anti-HSP70/HSC70 (ADI-SPA-797), fin. con. 1 µg/ml; and rabbit anti-calnexin (ADI-SPA-865), fin. con. 1 µg/ml. From Santa Cruz Biotechnology, we obtained rabbit anti-β1 integrin (M-106), fin. con. 2 µg/ml; goat anti-TSG-101 (M-19), fin. con. 2 µg/ml; and rabbit anti-CD63 (H-193), fin. con. 2 µg/ml. Mouse anti-PrP (6D11), fin. con. 0.5 µg/ml was obtained from Biolegend, mouse anti-CD9 (MM2/57), fin. con. 0.5 µg/ml was obtained from Merck; mouse anti-PrP (AH6), fin. con. 0.5 µg/ml was obtained from The Roslin Institute, and donkey anti-mouse Alkaline Phosphatase conjugate, fin. con. 0.3 µg/ml was obtained from Jackson ImmunoResearch.

### Cells

Prion infection models were used in the CAD5 cell line (originated from: RRID:CVCL_0199)[65], the N2a-PK1 cell line (RRID:CVCL_WI10)[66], and cells chronically infected with the RML5 prion strain (CAD5-RML, N2a-PK1-RML). The cells were maintained in Opti-MEM (Thermo Fisher Scientific) supplemented with 10% bovine growth serum (BGS, HyClone Bovine Growth Serum, GE Healthcare Life Sciences) and 100 U of penicillin/streptomycin/ml (Lonza). CAD5 cells were passaged by washing with 0.5 mM EDTA in PBS followed by resuspension in fresh medium. The N2a-PK1 cell line was passaged by washing with 0.5 mM EDTA in PBS and trypsinized (Trypsin-EDTA 0.25%, Thermo Fisher Scientific).

All relevant data on EV production, isolation, and characterization were submitted to the EV-TRACK knowledgebase (EV-TRACK ID: EV220318)[67]. The score for the CAD5 EVs were 67% and 56% for N2a-PK1 EVs. For EV production, the cells were passaged a day before production (∼ 30% confluence). Cells were washed with PBS (Lonza), and Exo-free Opti-MEM was added for conditioning with EVs. The exo-free Opti-MEM medium was produced as published [68]. Opti-MEM supplemented with 20% BGS and penicillin/streptomycin was centrifuged at 110,000 × g (SW28 rotor, Beckman) for 20 hours at 4 °C in an ultracentrifuge (LE-80K, Beckman). The centrifuged medium was diluted with serum-free Opti-MEM supplemented with penicillin/streptomycin to a final concentration of 10% BGS and filtered using the Stericup-VP Sterile Vacuum Filtration System with a 0.1 µM PES filter (Merck). The cells were incubated with the medium for 48 hours. When the conditioned medium was harvested, the cells had 100% confluence with a minimum of dead/floating cells.

### Isolation of EVs

EVs were isolated from the conditioned medium by a series of differential centrifugations. All steps were performed on ice, and samples were centrifuged at 4 °C. The conditioned medium was centrifuged at 300 × g for 15 min. The supernatant was collected and centrifuged at 2,000 × g for 20 min to sediment large cell debris. The supernatant was centrifuged at 10,500 rpm (∼20,000 × g, k factor = 1,753) for 70 min (SW28 rotor, Beckman). Pelleted lEVs were resuspended in 13.5 ml of PBS (Lonza) and sedimented again at 10,500 rpm (∼20,000 × g, k factor = 1,983) for 70 min (SW40 rotor, Beckman). The supernatant containing sEVs was centrifuged at 25,000 rpm (∼ 110,000 × g, k factor = 309) for 70 min (SW28 rotor). The pellet was resuspended in ∼ 12 ml of PBS and underlaid with 2 ml of 40% sucrose in 0.2 M Tris-HCl pH 7.4 and centrifuged under the same conditions. The sucrose cushion containing sEVs was collected with an 18 G needle, diluted with PBS to a final volume of 13.5 ml, and centrifuged again in SW40 rotor (k factor = 350). Both lEVs and sEVs were resuspended in PBS and frozen on dry ice. Pure EVs were used for western blot characterization and infection using fractions standardized to the same amount of protein (TPS) determined by the BCA assay (Pierce BCA assay kit, Thermo Fisher Scientific). The graphical scheme of isolation is shown in Figure 1.

**Figure 1:**
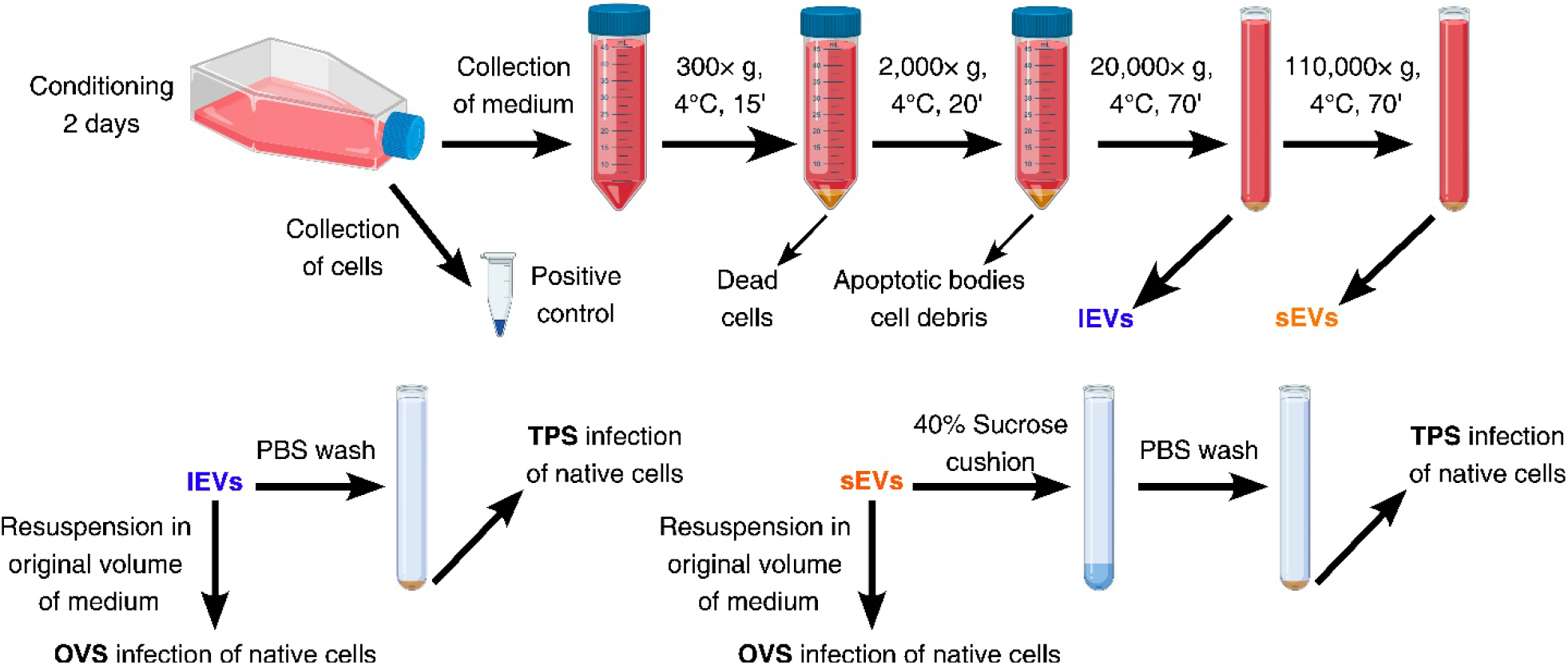
Scheme of lEV and sEV isolation and infection of native cells by the OVS and TPS. Created with BioRender.com

### Infection of cells

Native cells were passaged 24 h before infection. The cells were washed with 5 ml of PBS, and an Exo-free medium containing isolated EVs was added for ∼ 72 h. Two schemes of cell infection were used. In the first scheme, lEVs and sEVs were standardized according to the original volume of the collected sample. We abbreviated this scheme as OVS (Original Volume Standardization). The pelleted lEVs after the first 20,000 × g spin and sEVs after the first 110,000 × g spin were resuspended in a volume of Exo-free medium that was identical to the starting volume of the collected conditioned medium (8 ml). Aliquots of 4, 2, and 1 ml were used to infect native cells in a 25 cm^2^ flask or 100 µl in a 96-well plate. The second infection scheme was standardized to contain the same amount of total protein used to infect cells. We use TPS (Total Protein Standardization) as an abbreviation. Cells in a 25 cm^2^ flask were infected with 10, 5, or 2.5 µg of total protein in EV fractions. On a 96-well plate, the cells were infected with 0.2, 0.1, or 0.05 µg of EV fractions. The cells were passaged every 3-4 days to reach the 10^th^ or 12^th^ passage. Two independent experiments, including EV production and isolation, were carried out. As a positive control, the cell homogenates of RML-infected cells were used, and negative controls were homogenates of the noninfected cells. The graphical scheme of infection is shown in Figure 1.

### Harvesting of the cells after the infection for analysis

CAD5 or CAD5-RML cells were resuspended in 0.5 mM EDTA in PBS by pipetting. N2a-PK1 or N2a-PK1-RML cells were washed in 0.5 mM EDTA in PBS. The cells were centrifuged at 300 × g and 4 °C for 10 min, the supernatant was discarded, and the pellets were frozen at -80 °C. The pellets were thawed on ice, and freeze‒thaw cycles were performed 3 times in total. The cells were homogenized using a 30 G needle. To determine the protein concentration, part of the homogenate was lysed in lysis buffer (150 mM NaCl, 50 mM Tris-HCl pH 7.5, 0.5% (V/V) Triton X-100, 0.5% (w/V) sodium deoxycholate) along with part of the collected EVs. The samples were incubated on ice for 30 min with occasional vortexing and then centrifuged at 14,000 × g and 4 °C for 10 min. The supernatant was used to determine the protein concentration by BCA assay according to the manufacturer’s instructions (Pierce BCA Protein Assay Kit, Thermo Fisher Scientific).

### Cell blot

All steps were performed at room temperature unless stated otherwise. Infected cells were seeded in duplicate on a plastic coverslip in a 24-well plate and grown to confluency, as we published previously [69]. The coverslips were blotted on a 0.45 µm nitrocellulose membrane (Bio-Rad) wetted in lysis buffer. The membranes were dried and incubated in lysis buffer containing 7 µg/ml proteinase K (PK, Sigma‒ Aldrich) at 37 °C for 90 min. The membranes were washed 2× in Milli-Q H_2_O (MQH_2_O). The PK was stopped with 2 mM phenylmethylsulfonyl fluoride (PMSF) in PBS for 10 min and washed 3× in MQH_2_O. The membranes were denatured with 3 M guanidium thiocyanate in 10 mM Tris-HCl pH 7.5 for 10 min and washed with MQH_2_O 5×. After denaturation, membranes were blocked in 5% nonfat milk in 0.5% Tween in Tris buffer pH 7.4 (TBS-T) for 30 min. The primary antibodies anti-PrP 6D11 and AH6 were diluted in a blocking solution, and the membranes were incubated for 2 h at RT or overnight at 4 °C with antibodies. Membranes were washed with TBS-T 3× 5 min and incubated in secondary antibody conjugated to alkaline phosphatase diluted in blocking solution for 1 h. The membranes were washed with TBS 5× 5 min and developed using BCIP/NTP (Merck). The reaction was stopped using MQH_2_O. Densitometry was evaluated in AzureSpot software (v.2.0, Azure Biosystems) on background-corrected images.

### SDS‒PAGE and western blot

The frozen cell pellets were thawed on ice, resuspended in lysis buffer, and incubated on ice for 30 min with occasional vortexing. Lysed cells were centrifuged at 14,000 × g at 4 °C for 10 min. The collected supernatant was used to measure protein concentration using the BCA assay (Pierce BCA Protein Assay Kit, Thermo Fisher Scientific) according to the manufacturer’s instructions. The EVs were thawed on ice, and part was diluted in lysis buffer to measure protein concentration as in cell lysates. Nonlysed EVs were diluted with MQH_2_O to unify the protein concentration, and sample buffer was added (final concentration: 0.5 M Tris-HCl pH 6.8, 4% SDS, 0.125% bromophenol blue, 12.5% glycerol). Cell lysates were diluted with MQH_2_O to standardize the protein concentration before adding the sample buffer. To detect PrPres, lysates and EVs were incubated with PK (50 µg/ml) at 37 °C for 45 min prior to adding the sample buffer. The samples were boiled at 95 °C for 10 min. Boiled samples were loaded on 10% acrylamide/bisacrylamide gel (Bio-Rad) or TGX Stain-Free™ FastCast™ 12% acrylamide gel (Bio-Rad). Proteins on gels were imaged in an Azure c600 imaging system according to the manufacturer’s instructions. Separated proteins on the gels were blotted on Amersham™ Hybond® low-fluorescence 0.2 µm PVDF membranes (Merck). The membranes were blocked using fluorescence blocking buffer (Azure Biosystems) at RT for 30 min. Primary antibodies were diluted in blocking buffer and incubated at RT for 2 h or at 4 °C overnight on a rocker. The membranes were washed with IR washing buffer (Azure Biosystems) 3× 5 min. Washed membranes were incubated with secondary antibody diluted in blocking buffer at RT for 1 h. Secondary antibodies were washed 3× 5 min in IR washing buffer and 2× 5 min in PBS. After drying, the membranes were imaged using the Azure c600 imaging system. Densitometry was evaluated in AzureSpot 2.0 software using predefined 200-radius rolling ball background subtraction.

### Real-Time Quacking Induced Conversion (RT-QuIC)

The collected EV pellets were thawed on ice and homogenized with a 30 G needle, and the protein content was measured by BCA assay. Homogenized cells and EVs were diluted to 3.2 µg/µl using PBS, corresponding to 10% cell homogenate protein content. By decimal and quintuple dilutions, EVs and cell homogenate were diluted to 10^-6^ – 10^-9^ and 10^-5^ – 3×10^-11^, respectively, in PBS buffer containing 1× N-2 supplement (Gibco, Thermo Fisher Scientific) and 0.1% SDS and then analyzed by second-generation RT-QuIC assay. All samples were run in quadruplicate in a Nunc™ MicroWell™ 96-Well Optical-Bottom Plate (Thermo Fisher Scientific). Each well contained reaction mix (10 mM phosphate buffer, pH 7.4; 300 mM NaCl; 10 µM thioflavin T (ThT); 1 mM EDTA; 0.002% SDS; and 0.1 mg/ml rSHa PrP (90–231)) and 2 µL of sample in a volume of 100 µL. The reaction was carried out using the FLUOstar Omega plate reader (BMG LABTECH GmbH), undergoing repeating shaking cycles of 60 s (700 rpm, double orbital) and 60 s of rest for 60 h at 55 °C. Fluorescence was measured every 15 min. Each test was analyzed using Mars software (BMG LABTECH GmbH) [70,71].

### Standard Scrapie Cell Assay

All steps were performed at room temperature unless stated otherwise, as previously published [72]. Infected CAD5 cells were seeded in ELISPOT plates (multiscreen 96-well filter plates with high protein-binding Immobilon-P membrane, 0.45 µm, Millipore) activated with 70% ethanol. Ethanol was removed by vacuum (Supelco), and the wells were washed 2× with PBS. Cells (1,000, 5,000, or 10,000 per well) were transferred to the plates, immobilized on the membrane using a vacuum, and dried at 50 °C. The dried plates were incubated with 60 µl of PK (2 µg/µl) in lysis buffer at 37 °C for 30 min. PMSF (2 mM) was added in PBS to inhibit PK for 10 min. Cells were washed with PBS, and proteins were denatured with 3 M guanidium thiocyanate in 10 mM Tris-HCl pH 7.5 for 10 min and washed 4× with PBS. The membranes were blocked using Superblock™ solution (Thermo Fisher Scientific) for 1 h. After blocking, the membranes were incubated with 6D11 primary antibody diluted in Superblock™ solution at 4 °C overnight. The primary antibody was washed away 4× with TBS-T, and the membranes were incubated with a secondary antibody conjugated with alkaline phosphatase diluted in Superblock™ solution for 1 h. The secondary antibody was washed away 5× with TBS-T and developed with an AP Conjugate Substrate Kit (Bio-Rad). Spots were quantified using a Nikon ELISPOT system and NIS-Elements AR software (v. 4.20).

### Statistical Analysis

GraphPad Prism 5 (v. 5.03, GraphPad Software) was used for statistical analysis. Data were analyzed with the D’Agostino and Pearson omnibus normality test and the Shapiro–Wilk normality test. Two-tailed parametric unpaired t-test was used where the data passed the normality test (Standard Scrapie Cell Assay). Two-tailed Wilcoxon matched-pairs signed rank test (non-parametric test) was used where the data did not pass the normality test (Cell blots and Western blots). The paring corresponded with the used infection dose of lEV and sEV infection. In the case of Western blot, the pairing also corresponded with the protein loading on the gel. All the data are presented as the mean with the standard error of the mean (SEM). Each spot on the presented graph represents a biological or technical replicate of infection.

### Electron Microscopy

For transmission electron microscopy (TEM), EM grids (FCF200-Cu, Electron Microscopy Sciences) were prepared as published [68,73]. Isolated EVs were diluted 10× in PBS. Grids were incubated for 20 min on a 15 µl drop of EV suspension and transferred to a 50 µl drop of 4% formaldehyde in PBS for another 20 min. The fixed samples were washed on a 100 µl drop of PBS and fixed again on a 50 µl drop of 1% glutaraldehyde in PBS (electron microscopy grade, Sigma‒Aldrich) for 5 min. The grids were washed 8× on a 100 µl drop of MQH_2_O. Washed grids were contrasted on 25 µl drops of uranyl oxalate pH 7 for 5 min. Contrasted grids were transferred on 25 µl drops on ice of a mix of 4% uranyl acetate and 2% methylcellulose (1:9) for 10 min. The mix was slowly removed with filter paper, and the grid was air-dried. Cryo-TEM samples were prepared by plunge freezing [73,74]. The EV sample (3 µl) was applied to Lacey carbon grids (LC-200 Cu, Electron Microscopy Sciences) covered with a perforated carbon film and discharged for 40 s with a 5 mA current. Most of the sample was removed by blotting (∼ 1 s), and the sample was immediately plunged into liquid ethane kept at -183 °C. The grids were transferred without reheating to a Gatan 626 cryo-specimen holder and imaged in a Tecnai G2 Sphera 20 transmission electron microscope equipped with a LaB6 gun and a Gatan UltraScan 1000 slow-scan CCD camera. The images were recorded at an accelerating voltage of 120 kV. All pictures were CTF-corrected and bandpass filtered to suppress ice thickness and noise below 1 nm detail size.

## Results

### Characterization of lEV and sEV fractions

The isolated fractions of lEVs and sEVs from CAD5-RML and N2a-PK1-RML cells differed in protein composition (Figures 2A and B). Common markers for EVs, Alix, HSP70, TSG-101, CD63, and CD9, were present in both fractions of EVs. In contrast, we did not detect calnexin, a marker of contamination by the endoplasmic reticulum, in our EV fractions. In the CAD5-RML lEV fraction, very weak bands were detected by calnexin antibody (Figure 2C, Supplementary Table S1). To distinguish lEVs from sEVs, we used β-1 integrin as a marker (Figure 2C, Supplementary Table S1). Large EVs, which should consist mainly of MVs generated by budding from the plasma membrane, should be enriched in this marker compared to sEVs. In CAD5-RML cell line, we isolated sEVs depleted of β-1 integrin (Figure 2C, Supplementary Table S1), but sEVs from N2a-PK1-RML cell line were not completely depleted of β-1 integrin (Figure 2D, Supplementary Table S1). The sEV fractions were enriched in the tetraspanin markers CD9 and CD63, TSG-101, and Alix compared to the lEVs in both cell lines. The exact densities are summarized in Supplementary Table S1. The presence of EVs in isolated fractions was verified by electron microscopy. As shown in Figures 2E and F, CAD5 lEVs and sEVs differed in size. The size difference was also visible in the N2a-PK1 cell line (Figures 2G and H), although in sEVs, the vesicles were larger compared to CAD5 cell sEVs. In addition, we used cryo-electron microscopy to visualize the ultrastructure of CAD5 lEVs and sEVs where lipid bilayers were visible (Figure 2I-L) and to measure the vesicles. We found that the mean size (n = 200) of sEVs was 79 nm (SEM ±3 nm, Supplementary Figure S1). The largest vesicles in the sEV fraction were approximately 160 nm in diameter. The sizes of the visible vesicles in the lEV fraction ranged from ∼100 nm – ∼500 nm. Larger vesicles were present but deformed due to the thickness of the ice or holes in the perforated carbon film (Figure 2I).

**Figure 2:**
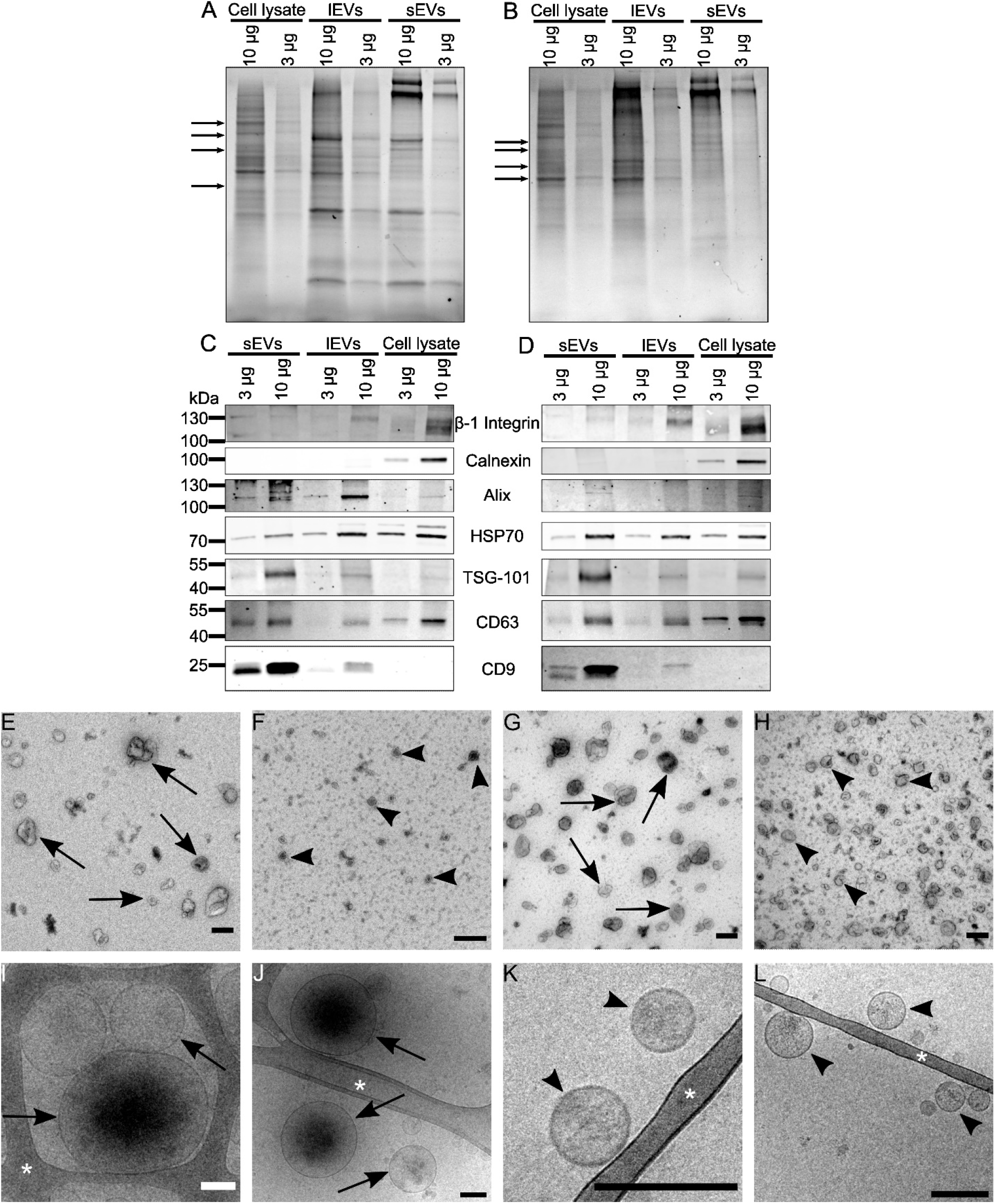
Characterization of lEV and sEV fractions. (a,b) An example images of the TGX Stain-Free™ SDS–PAGE protein profile of CAD5-RML (a) or N2a-PK1 (b) cell lysate, lEVs, and sEVs with the loading of 3 µg or 10 µg of proteins in line. Arrows point to examples of protein profile differences between cell lysate, lEVs, and sEVs. (c,d) An example of images of the detection of markers in the EV fraction from the CAD5-RML (c) or N2a-PK1-RML (d) cell line. β-1 integrin: marker of lEVs; calnexin: endoplasmic reticulum marker – contamination of EV fractions; Alix, HSP-70, TSG-101, CD63 and CD9: common markers for sEVs and lEVs. (e-g) An example images of EVs present in the lEV (e, g) and sEV (f, h) fractions isolated from CAD5 (e, f) or N2a-PK1 (g, h) cells contrasted with uranyl oxalate. Bars: 200 nm. (i-l) Cryo-electron microscopy images of vesicles in lEV (i, j) and sEV (k, l) fractions from CAD5 cells. Arrows: EVs in lEVs fraction, Arrowheads: EVs in sEVs fraction, Asterix: Carbon Lacey support film, Bars: 200 nm. For visualization purposes, all images were enhanced with brightness and contrast.

### Large and small EVs are enriched in PrPC and PrPres content compared to intact cells

We found that PrP in both EV fractions was significantly enriched compared to the whole-cell lysates of corresponding cell lines (Figures 3 A-D). When PrPres was normalized to whole-cell lysate PrP PK-(PrP^C^ + PrP^Sc^), we found that in lEVs, there was approximately five times more PrPres in CAD5-RML cells and approximately 18 times more PrPres in N2a-PK1-RML cells. In sEVs, there were approximately 8 and 11 times more PrPres than in the cell lysates of CAD5-RML cells and N2a-PK1-RML cells, respectively. There was approximately 5× less PrPres in whole-cell lysate compared to PrP PK-in CAD5-RML cells and approximately 20× less in N2a-PK1 cells. While the amount of PrPres was significantly 1.5× higher in lEVs than in sEVs from N2a-PK1-RML cells, the amount of PrPres in lEVs from CAD5-RML cells was slightly lower than that in sEVs (Figure 3E). Similar patterns were also recorded for the amounts of PrP PK-in the EV fractions in the CAD5-RML. Large EVs had also 1.5× higher amount of PrP PK-than sEVs (Figure 3F).

**Figure 3:**
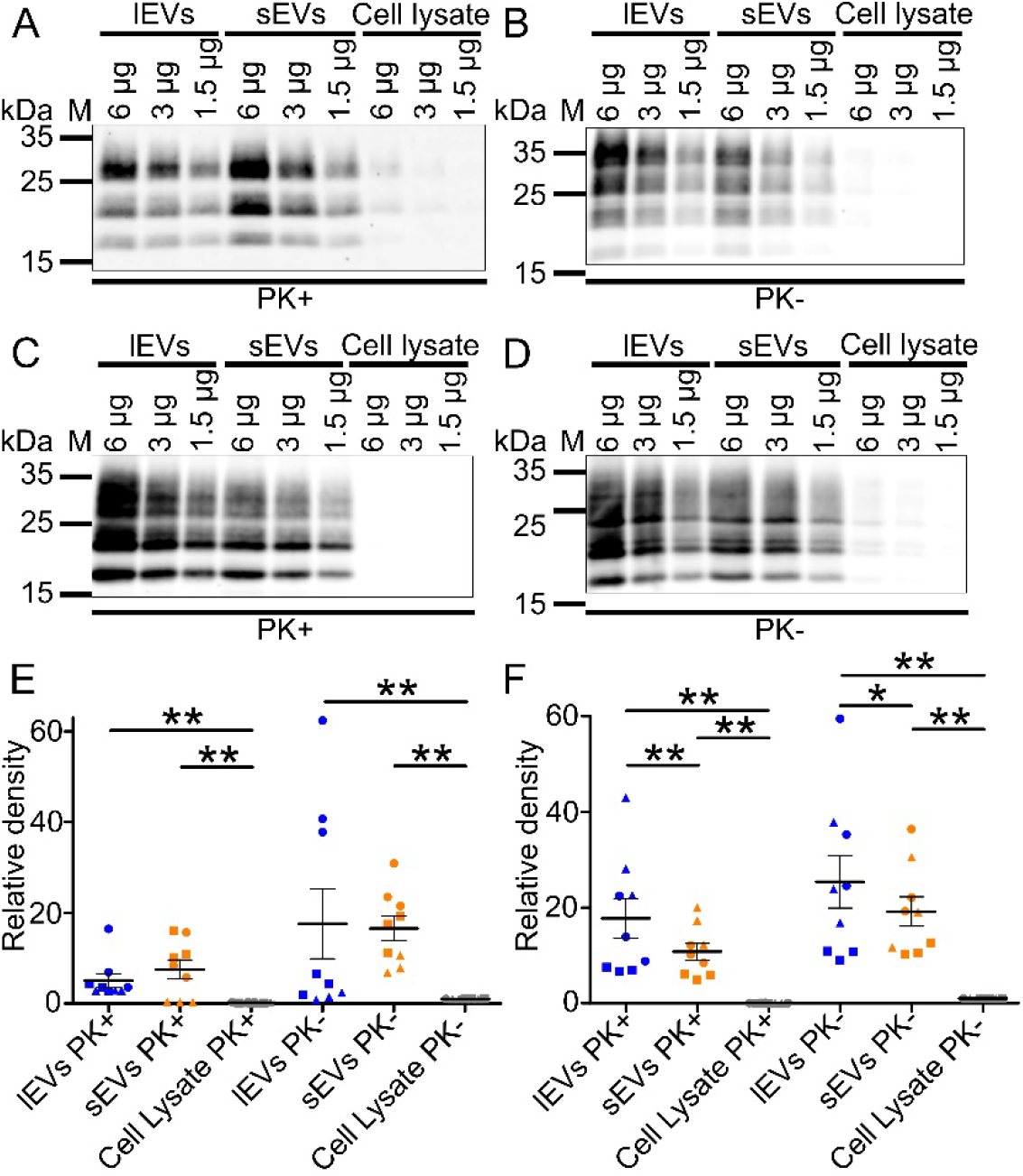
western blot analysis of PrP levels in EV fractions. (a-d) CAD5-RML (a, b) or N2a-PK1-RML (c, d) lEVs, sEVs, and whole-cell lysate were treated with proteinase K (PK+, a, c), and PrPres was detected using a mix of prion antibodies 6D11 and AH6. Untreated (PK-) samples (b, d) demonstrated the presence of total PrP (PrP^C^ + PrP^Sc^). For visualization purposes, all images were enhanced with brightness and contrast. (e,f) Comparison of the relative densities of PrPres bands in lEVs, sEVs, and whole-cell lysate normalized to PrP PK-signal in whole-cell lysate from CAD5-RML (e) or N2a-PK1-RML (f) cells. The line represents the mean with SEM of biological triplicates, each with technical triplicates (3 different protein loadings on gel – 6, 3, or 1.5 µg per line). Since the relative densities are normalized to whole-cell lysate which was present on all of the membranes, the densities in dot plots are comparable. Blue spots – lEVs, orange spots – sEVs. Circles, squares, and triangles are marking individual biological replicates of isolation. Samples were analyzed by two-tailed non-parametric paired t-test (Wilcoxon matched-pairs signed rank test) with pairing of corresponding protein loading. * P < 0.05; ** P < 0.01.

### Large EVs have higher prion-converting activity than small EVs

We used an endpoint dilution RT-QuIC assay to determine the prion-converting activity in the EV fractions standardized to contain the same amount of total protein. The seeding dose 50 (SD_50_) of CAD5-RML lEVs was twenty times higher than that of sEVs (2×10^-8^ vs. 4×10^-7^, Figure 4E). Additionally, the maximum relative fluorescence at a 10^-5^ dilution was higher and the lag phase shorter in lEVs (163,000 and 6.5 h, Figure 4A) than in sEVs (151,000 and 8 h min, Figure 4B). In N2a-PK1-RML cells, the difference was less convincing. In the first isolation of EVs, there was approximately 26× higher prion-converting activity in lEVs (SD_50_: 3×10^-9^ vs. 8×10^-8^) and the maximum relative fluorescence and lag phase of the 10^-5^ dilution were 184,000 and 9.5 h for lEVs and 119,000 and 14.3 h for sEVs, respectively (Figure 4C and D). In the second isolation, no difference was recorded (SD_50_: 3×10^-9^ vs. 3×10^-9^) and the respective values were 152,000 and 12.3 h in lEVs and 146,000 and 11.5 h in sEVs of the 10^-5^ dilution. The numbers of positive wells in quadruplicate for each sample and dilution are summarized in Supplementary Table S2.

**Figure 4:**
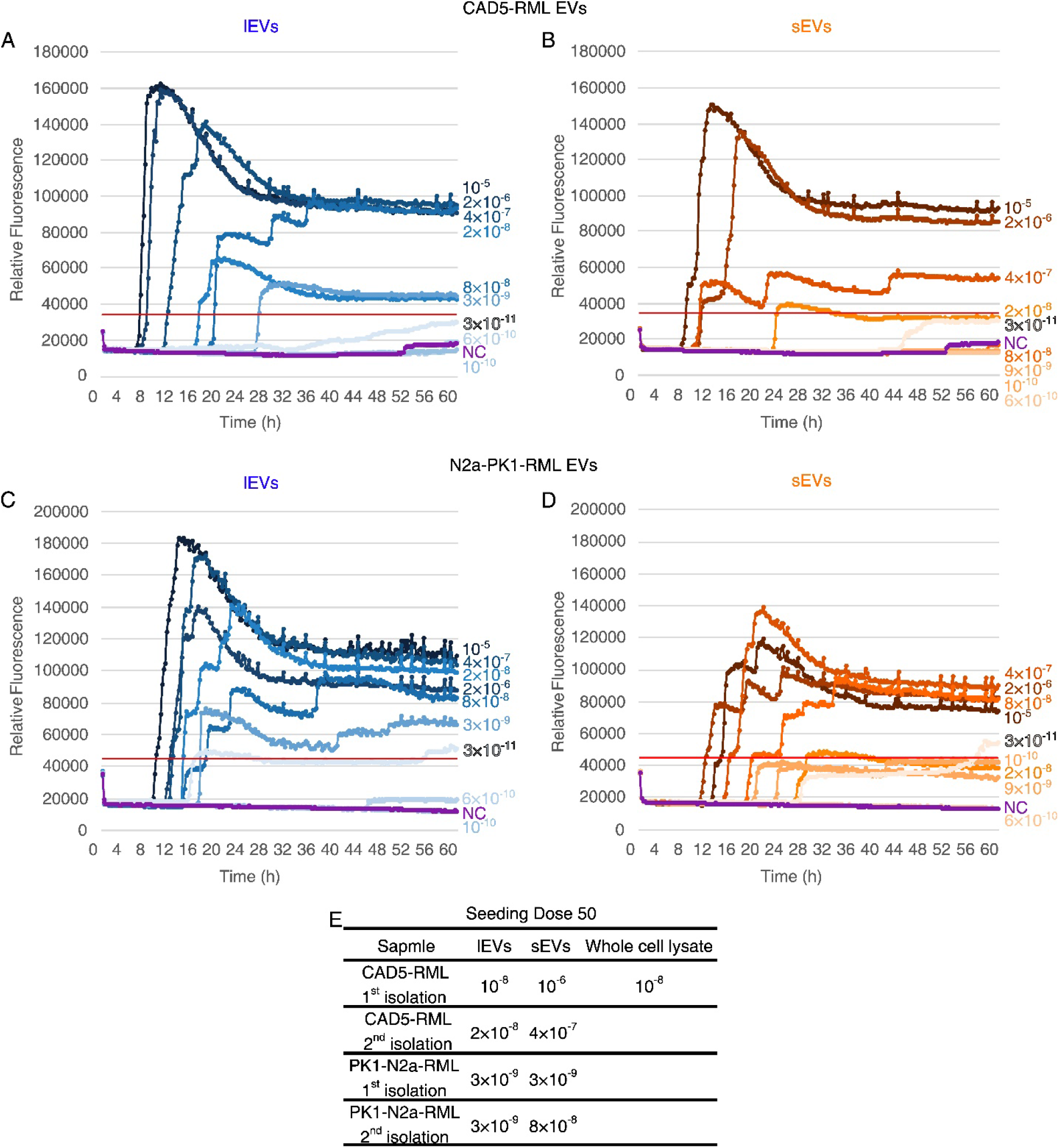
Comparison of EV fractions’ prion converting activity using endpoint dilution RT-QuIC assay. (a-d) Quintuple dilution series of CAD5-RML lEVs (a), CAD5-RML sEVs (b), N2a-PK1-RML lEVs (c), and N2a-PK1-RML sEVs (d). Each line in the graphs represents the mean value of the technical quadruplicate. Redline – threshold (mean plus 5× standard deviation from negative control). NC (purple line) – negative control (uninfected CAD5 or N2a-PK1 whole cell lysate at 10^-5^ dilution, corresponding to 0.001% cell homogenate). (e) Table with seeding dose 50 from two independent isolations of EVs. The value represents the highest dilution, where at least 2 out of 4 technical replicates gave positive results. The whole cell lysate SD50 was determined only in the first isolation from CAD5-RML cells. In other three isolation, whole cell lysate served only as a positive control.

### A fraction of lEVs is more effective in the infection of cultured cells than a fraction of sEVs

After fraction characterization, we infected native CAD5 or N2a-PK1 cells with EVs isolated from the corresponding chronically infected cells. First, we infected native CAD5 cells with supernatants collected during EV isolation (Supplementary Figure S2) to monitor the loss of infectivity produced by individual centrifugation steps. Most of the infectivity was lost after pelleting floating cells (300× g). The next visible loss of infectivity occurred after pelleting of the lEVs. Some infectivity remained in the conditioned medium even after pelleting sEVs (110,000× g supernatant) as demonstrated by visible infection of the cells after 10 passages (Supplementary Figure S2F). The results from supernatant infection were in agreement with the infection with collected pellets during the isolation procedure (Supplementary Figure S3). We compared cell blots of infected cells with lEVs and sEVs in passages 3, 5, 8, and 10 (Figure 5A and B). We found that lEVs infected native CAD5 cells more effectively than sEVs and verified the results on N2a-PK1 although with lesser differences between EV fractions. We used two schemes of infection: an OVS (original volume standardization) reflects the total amount of prion infectivity associated with the isolated EV fractions, and a TPS (total protein standardization) reflects the concentration of prion infectivity present in the isolated EV fractions. The differences in cell blot by OVS infections were uniform during the passages after infection of CAD5 cells. Infection by lEVs was significantly 1.5× - 2.3× stronger - 1.7× (p = 0.0147), 1.5× (p = 0.0016), 1.5× (p = 0.0165) and 2.3× (p > 0.05) in passages 3, 5, 8, and 10, respectively, Figures 5C and D). Similarly, infection by lEVs was significantly stronger in TPS infection. In earlier passages, the difference was smaller. In passage 3, lEV infection was 1.4× stronger (p > 0.05), and in passage 5, it was 1.6× stronger (p = 0.0018). In later passages 8 and 10, the infection was 4.1× stronger (p < 0.0001 and p = 0.0004), Figures 5C – D). In the N2a-PK1 cell model, significantly stronger infection by lEVs was confirmed. The OVS led to 1.4× - 1.7× stronger infection by lEVs (1.7× (p > 0.05), 1.4× (p = 0.0488), 1.5× (p = 0.0005) and 1.4× (p = 0.0008) in passages 3, 5, 8, and 10, Supplementary Figure S4). In the TPS scheme of infection, lEVs infected N2a-PK1 cells 1.2× significantly strongly in passages 8 and 10 (p = 0.0065 and 0.0007. In passage 3, the results were not significant and in passage 5, sEVs infected native cells significantly 1.2× stronger (p = 0.0021)(Supplementary Figures 6C and D).

**Figure 5:**
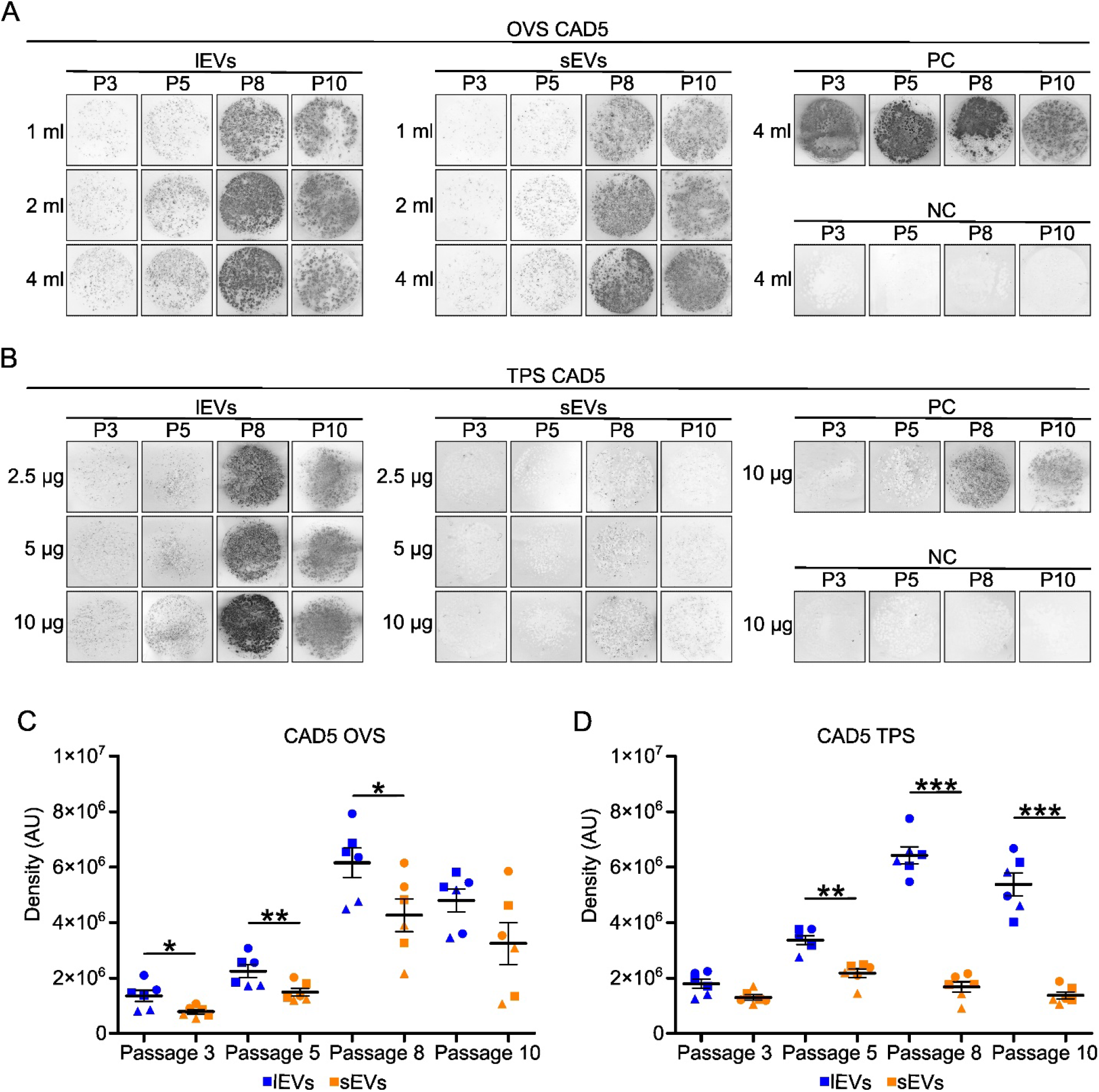
Example images of cell blots from passages after infection of CAD5 cells with lEVs and sEVs with densitometry analysis. (a and b) Example images of cell blots from infection of native CAD5 cells infected with lEVs and sEVs in different passages after infection (passage 3, 5, 8, and 10). Cells were infected with 4 ml, 2 ml, and 1 ml of resuspended EVs in OVS (a) infection or with 10 µg, 5 µg, and 2.5 µg of EVs in TPS infection (b). NC – negative control, PC – positive control. For visualization purposes, all images were enhanced with brightness and contrast. (c and d) Densitometry analysis of cell blots from different passages. Infection with lEVs was compared to the corresponding infection with sEVs. The line represents the mean with SEM. Blue spots – lEVs, orange spots – sEVs, circles – infection with 4 ml or 10 µg, squares - infection with 2 ml or 5 µg, triangles - infection with 1 ml or 2.5 µg. Each spot is average of technical duplicate. Samples were analyzed by two-tailed non-parametric paired t-test (Wilcoxon matched-pairs signed rank test) with pairing of corresponding infection dose. * P < 0.05; ** P < 0.01, *** P < 0.001. AU – arbitrary units.

After cell blot analysis, we confirmed the results with western blotting from each passage and infection (Figure 6 and Supplementary Figures S5 – S7). From the same samples, western blot analysis showed that infection of CAD5 cells by lEVs was even stronger compared to cell blot analysis. With the OVS scheme, the PrPres signal from lEV-infected CAD5 cells was 1.5× - 3.6× significantly stronger then from sEV-infected CAD5 cells (1.5×, 3.6×, 3.2×, and 1.9× in passages 3, 5, 8, and 10, Figure 6E and Supplementary Figures S5 - 7). TPS infection by lEVs in CAD5 cells led to a much higher PrPres signal in the CAD5 cell model than infection with sEVs. The PrPres signal reached its maximum (where the dilutions had minimal differences) in passage 8, but infection by sEVs just started to appear (Figure 6C). Within these limitations, we measured a 24.6× significantly stronger PrPres signal in passage 8. Since lEV infection reached its maximum and infection by sEVs continued to increase, the difference dropped to 7.0× and 10.6× in passages 10 and 12, respectively (Supplementary Figure S7E). In the N2a-PK1 cell line, the trend of stronger infection by lEVs was similar. Unlike CAD5 cells, differences in TPS infection were smaller than in OVS infection in N2a-PK1 cells but remained significant. TPS infection led to 1.4× - 1.8× stronger PrPres signals in lEV-infected cells than in sEV-infected cells (Figure 6E and Supplementary Figures S5 - 7). In earlier passages of the OVS scheme of infection, the PrPres signal was 2.0× and 1.9× stronger in infections by lEVs in passages 3 and 5, respectively (Supplementary Figures S5 – S7). Later passages after infection by lEVs revealed a 4.8× significantly stronger PrPres signal in passage 8 and a 3.0× stronger signal in passage 10 (Figure 6E and Supplementary Figure S7). The absolute density values for each sample are summarized in Supplementary Figures S9 and S10. As controls, we compared the total PrP signal of non-digested PrP PK-samples in the same manner as PrPres (Figure 6F). Despite the different strengths of infection by lEVs and sEVs, cells infected by both types of EVs retained similar levels of PrP PK-. The lowest PrP PK-the ratio of lEVs/sEVs (0.96×) was in passage 5 with OVS of CAD5 cells (Supplementary Figure S6F). The highest ratio of 1.84× was in the 10^th^ passage with OVS of N2a-PK1 cells (Supplementary Figure S6F). The only significant differences were at passage 10 in the infection of N2a-PK1 cells with both infection schemes. In both cases, lEV infected cells had higher levels of PrP PK-. As a loading control for PrP PK-, β-actin was used (Supplementary Figure 8). Relative densities of β-actin bands were similar or higher in samples infected with sEVs than with lEVs, confirming that the higher PrPres signal was not caused by elevated protein load.

**Figure 6:**
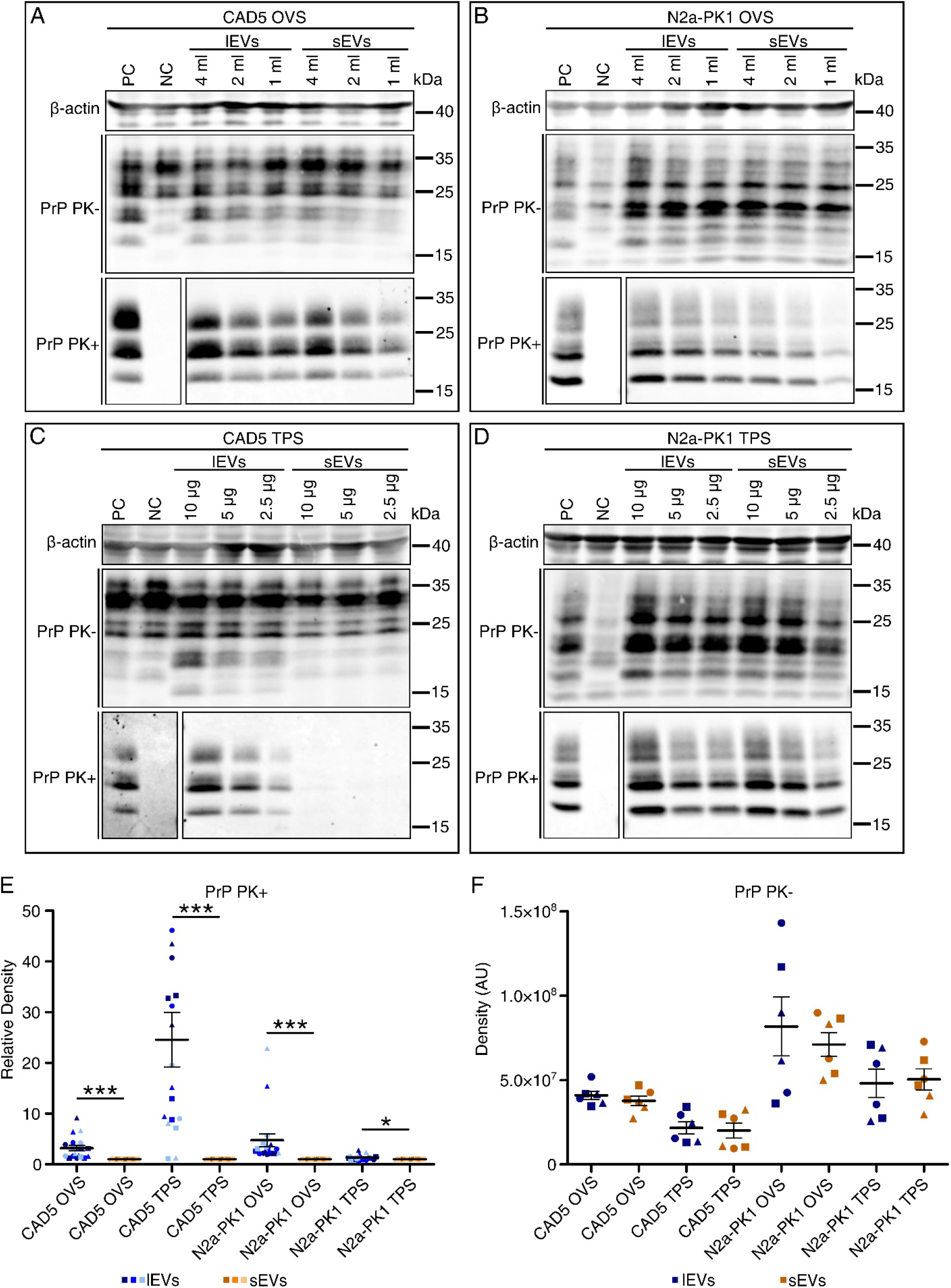
western blot analysis of PrPres and PrP^Sc^/PrP^C^signals in cells infected by lEV and sEV fractions at passage 8. (a-d) An example images of western blots from infection of CAD5 (a, c) or N2a-PK1 (b, d) cells with lEVs and sEVs at passage 8 after infection. Cells were infected with 4 ml, 2 ml, and 1 ml of resuspended EVs in OVS infection (a, b) or 10 µg, 5 µg, and 2.5 µg of washed Evs in TPS infection (c, d). PK+ - proteinase K digested samples. PK- - proteinase K untreated samples. β-Actin was used as a loading control for PrP PK-samples, which were also compared. All samples are shown for a loading of 40 µg per line. NC – negative control (mock-infected cells), PC – positive control. For visualization purposes, all images were enhanced with brightness and contrast. (e) Relative comparison of PrP signal in PrP PK+ samples by densitometry. The line represents the mean with SEM. Blue colors – lEVs, orange colors – sEVs. Light blue/orange – loading 10 µg per line, blue/orange – loading 20 µg per line, dark blue/orange – loading 40 µg per line. Circles – infection with 4 ml or 10 µg, squares – infection with 2 ml or 5 µg, triangles – infection with 1 ml or 2.5 µg. Samples were analyzed by two-tailed non-parametric paired t-test (Wilcoxon matched-pairs signed rank test) with the pairing of corresponding infection dose and protein loading per line. * P < 0.05; ** P < 0.01, *** P < 0.001. The quantification precision is limited in the CAD5 TPS infection – the PrPres signal in infection by lEVs was reaching the maximum while the PrPres signal started to appear in sEV infected cells. (f) Comparison of PrP PK-bands density. Blue spots – lEV infected cells, orange spots – sEV infected cells, circles – infection dose 4 ml or 10 µg, squares - infection dose 2 ml or 5 µg, triangles - infection dose 4 ml or 10 µg. The same statistical approach was used for PrP PK-analysis. AU – arbitrary units.

As another validation method, we used the standard scrapie cell assay (SSCA) to quantify infection for CAD5 cells precisely. The infections were carried out in hexaplicates for two independent EV isolations (Figure 7 and Supplementary Figure S11). In SSCA, dilutions in the TPS scheme reached their maximum level of infection in passage 8. We found a significant difference in passage 5. The mean spot counts (± SEM) in the 0.2 µg infection were 250 ± 20 spots for lEVs and 188 ± 18 spots for sEVs (p = 0.0309). At 0.1 µg, there were 206 ± 23 spots for lEVs and 113 ± 12 spots for sEVs (p = 0.0016) (Figure 7B). The differences were insignificant in passages 3 (Supplementary Figure S11B) and 8 (Supplementary Figure S11F). The mean spot counts (± SEM) for IEVs in passage 8 were 553 ± 18, 520 ± 16, and 353 ±34 for 0.2, 0.1, and 0.05 µg of total protein, respectively. In sEVs, infection yielded 452 ± 48 spots in 0.2 µg, 433 ± 37 in 0.1 µg, and 322 ± 52 in 0.05 µg infection. In the OVS infection, we found significant differences in passage 3 (Supplementary Figure S7D) and passage 8 (Figure 7D) but not in passage 5 (Supplementary Figure S7H). The mean spot counts were 511 ± 56 for lEVs and 267 ± 69 for sEVs (p = 0.0054, Figure 7D) for passage 8. In passage 3, the mean spot counts were 163 ± 17 for lEVs and 76 ± 12 for sEVs (p = 0.0004, Supplementary Figure S7D). All counted spots are summarized as the mean ± SEM in the Figure 7E).

**Figure 7:**
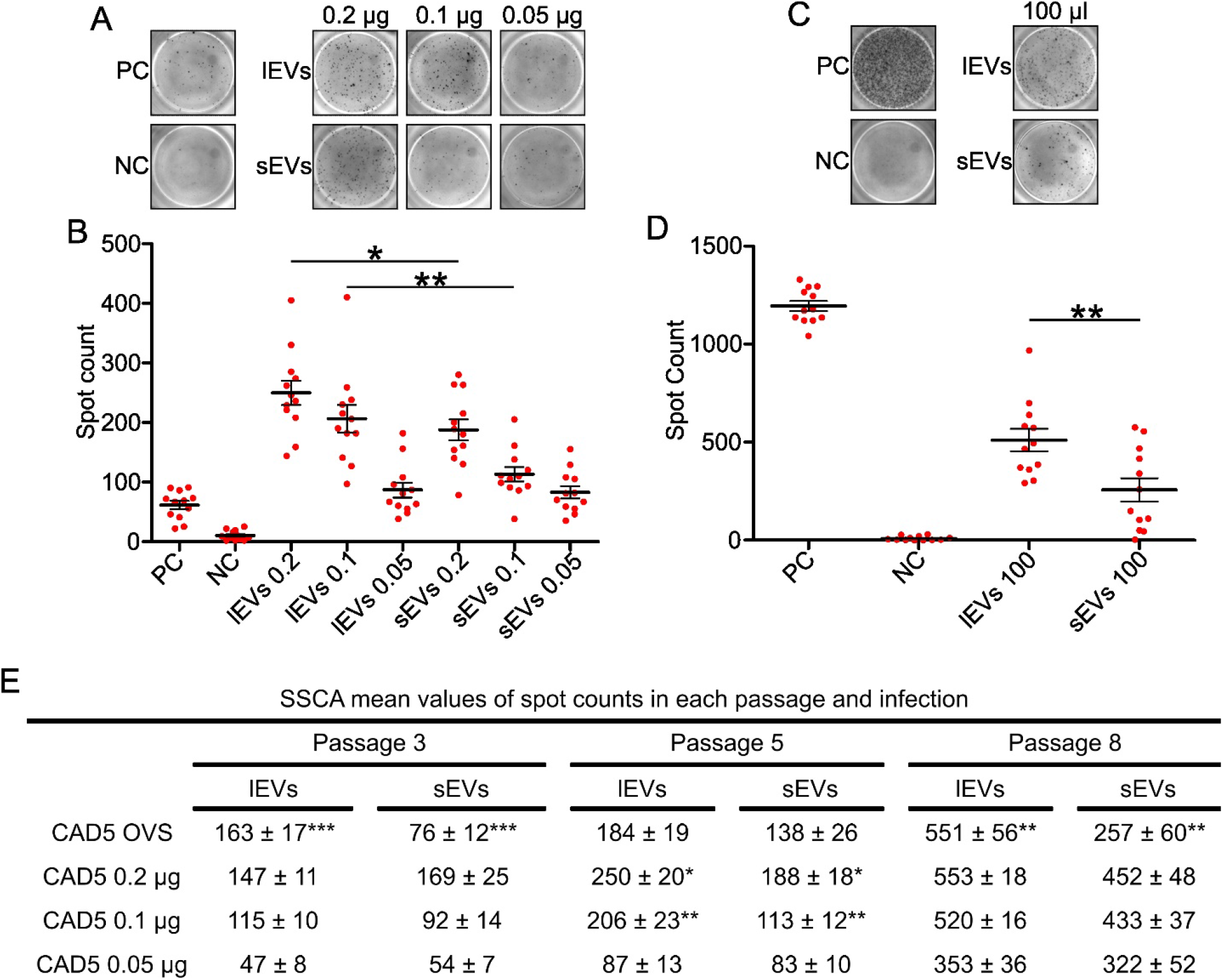
Scrapie cell assay of CAD5 cells infected with EVs. (a) Example images of wells in the scrapie cell assay infected with the TPS scheme in passage 5. PC – positive control (infection by CAD5-RML cell homogenate), NC – negative control (infection by CAD5 cell homogenate). A total of 1000 cells were transferred to each well. (c) Representative images of wells in the scrapie cell assay infected with the OVS scheme in passages 8. A total of 10,000 cells were transferred to each well. For visualization purposes, all images were enhanced with brightness and contrast. (b) Spot counts in each well of the scrapie cell assay (hexaplicates of infection from a duplicate of isolation) of TPS infection in passage 5. Each circle represents the count in one well. (d) Spot counts in each well of the scrapie cell assay in OVS infection at passage 8. The thick line represents the mean value with SEM, statistical significance was determined using a two-tailed unpaired t-test. (e) Table with summarized mean values ± SEM for infection by lEVs and sEVs in all passages and infection doses. ***P < 0.001 **P < 0.01 *P < 0.05.

## Discussion

The presence of infectious prion protein in exosomes was demonstrated by Février et al. [14], which is consistent with cellular prion conversion sites in the endolysosomal compartment, including MVBs [49,50]. However, one of the conversion sites of prion protein is the plasma membrane [55], and Mattei et al. [58] found PrP^TSE^ in microvesicles. PrP^TSE^ can be transmitted via EVs in cell culture [52]. According to Godsave et al. [75], approximately 50% of PrP^TSE^ is on the cell surface, 30% in invaginations, and only approximately 20% in cellular vesicles in the infected hippocampus. Although the predominant location of PrP^TSE^ is on the cellular surface, no one has compared microvesicles and exosomes in the spread of infection. To address this issue, we used a CAD5 cell model of prion infection infected with the RML prion strain. As a second cell line, we chose N2a-PK1 cells. Although both cell lines were selected for high prion production, CAD5 was produced by targeted oncogenesis of mouse catecholaminergic cells [76], and N2a-PK1 was derived from mouse neuroblastoma cells [66]. We tested the isolation protocol for microvesicles and exosomes to collect enriched fractions of these vesicles. According to the International Society of Extracellular Vesicles (ISEV) nomenclature and its position paper [64], we describe our collected fractions as large EVs, pelleted at 20,000 × g and enriched in β-1 integrin, possibly predominated by microvesicles. The sEVs were pelleted at 110,000 × g and enriched in CD9, CD63, TSG-101, and Alix. This fraction should be predominated by exosomes. Our cryo-EM experiments identified lEV and sEV fractions overlap in size. However, MVs and exosomes overlap in size according to several reviews defining EV sizes [77,78] and we cannot define MVs and exosome only by size. Although we optimized the isolation procedure, we could not isolate the pure sEV fraction without β-1 integrin, which is one of the markers for microvesicles [35], but it was recently also found in exosomes as another possible microvesicle marker [36]. This would explain the presence of β-1 integrin in our sEV fraction. Recently, Annexin A1 was found to be a specific marker for microvesicles [36]. We found enrichment of CD9 tetraspanin in sEVs compared to lEVs, as published previously [79]. Based on the detection of β-1 integrin in the lEV fraction, enrichment of CD9 in the sEV fraction compared to the lEV fraction, and no calnexin contamination, we believe that our lEV fraction is predominated by microvesicles and the sEV fraction by exosomes.

When we compared the EV fraction to the whole-cell lysate, we found great enrichment of both PrP^TSE^ and PrP^C^. The fraction of lEVs from CAD5 cells could not be isolated without contamination by calnexin under the various centrifugations we tried during isolation optimization. Typically, the presence of calnexin signals contamination from other cellular compartments [64]. In our lEV and sEV fractions, there was not a 100 kDa band of calnexin but three somewhat smaller bands that appeared in cells during apoptosis [80], visible only after immense overexposure of the membrane (three times longer exposure of the membrane or stretching of the histogram from 16 bit to 12 bit). Therefore, we assume that apoptotic bodies could cause this contamination since they overlap in size.

Isolated fractions of EVs could be normalized in two possible ways. By total lipid concentration [81] or total protein concentrations. Since we looked at prion protein transmission, we normalized EVs to protein concentration using the BCA assay. Measurement of particle number in two fractions distinct in size, which are heterogeneous, is problematic. Counting particles of different sizes by resistive pulse sensing (RPS) did not yield reliable results. A fixed pore size is needed for reproducible measurement, yet lEVs require a large pore size [82], with which sEVs are not detected [83]. For NTA, this method has a high relative concentration error for beads of 400 nm [84]. The minimum sizes for measuring vesicles are 70-90 nm by NTA and 70-100 nm by RPS [84]. In our sEV fraction, we detected the smallest particles with a size of 30 nm by cryo-TEM. Furthermore, processing software may induce artifacts since the two fractions need different settings [84].

We isolated both fractions from the same source, a conditioned medium, allowing us to compare these fractions in PrP levels directly. The PrP PK-levels in lEVs and sEVs were relatively the same in the CAD5-RML model, but in the N2a-PK1-RML model, the PrP PK-levels were slightly higher in lEVs. When we looked at PrPres levels in CAD5 EVs, they were higher in sEVs. The difference in N2a-PK1 EVs was in the opposite direction; the levels of PrPres were higher in lEVs. Our RT-QuIC experiments revealed higher conversion activity in lEVs than in sEVs from CAD5-RML EVs. However, in EVs from N2a-PK1-RML cells the prion converting activity was higher in one EV isolation but in second, the converting activity was the same in both fractions.

Next, we examined the infectivity of EVs by infecting native cells. First, we collected supernatants and pellets from all steps of the isolation protocol to map the distribution of prion infectivity in the CAD5 model. As expected, all the fractions contained the infectivity, including the fractions of isolated EVs. In accordance with their higher RT-QuIC conversion activity, the lEVs caused approximately four times higher CAD5 cell infection than sEVs in our initial cell blot experiments. We specifically looked at the difference between EVs by two distinct approaches. The OVS scheme aimed to evaluate the differences in the production of EVs by cells. Large EVs and sEVs were isolated from the same number of cells, and we omitted the washing steps to eliminate the loss of EVs as much as possible. The TPS scheme aimed to explore differences in the concentration of prion infectivity in washed vesicles and possibly also in their uptake by the cells. With both infection schemes, we obtained concurrent results demonstrating higher infectivity of lEVs in the CAD5 model. To overcome the limitations of cell blotting developed with alkaline phosphatase, which is prone to signal saturation, we utilized fluorescent western blotting with greater linear range of signal detection. The differences were even higher than in the cell blot. To further demonstrate stronger infection by lEVs, we utilized SSCA to count number of infected cells directly. Interestingly, in SSCA, we reached the infection plateau earlier (between the 5th and 8th passages) compared to the cell blot and western blot (8th – 10th passages). In the fifth passage, we had significantly more spots in TPS infection by lEVs. In OVS infection, we had significantly more spots in the 3rd and 8th passages in infection by lEVs. The differences measured by SSCA were less dramatic, but consistent with the results of the cell and western blot analysis.

To confirm stronger infection by lEVs in the CAD5 model, we similarly tested the infection by EVs in the N2a-PK1 cell line. Infection by lEVs was also stronger than that by sEVs with both infection schemes. In CAD5 models, it is interesting to note that the TPS scheme infection led to a greater difference between lEVs and sEVs suggesting higher concentration of the infectivity in lEVs. However, in the N2a-PK1 model, the TPS infection scheme led only to small difference suggesting similar concentration of infectivity in large and small EVs. In comparison, OVS infection scheme resulted in larger differences indicating enhanced production of lEVs by N2a-PK1 cells. To purify sEVs from aggregated PrP (also pelleted at 110,000 × g), we introduced a sucrose cushion in the isolation of sEVs [85]. Without the sucrose cushion step, the measurement would be biased by pelleted PrP aggregates. The effectiveness of the cells infection by lEVs and sEVs could be affected also by differences in their uptake. Since it is problematic to separate microvesicles and exosomes, there is limited knowledge of the uptake of different fractions of EVs. EVs can function as ligands for the surface receptor or be internalized by phagocytosis, macropinocytosis, caveolin-mediated endocytosis, clathrin-mediated endocytosis, lipid raft-mediated endocytosis, or fusion with the plasma membrane [32].

The CAD5 cells infected with the OVS scheme had a greater difference between infection levels in lEVs and sEVs compared to the TPS scheme. Conversely, in N2a-PK1 cells, the TPS scheme resulted in a higher discrepancy between infection levels in lEVs and sEVs compared to the OVS scheme. One possible explanation could be in the cell origin. CAD5 is derived from Cath.a cells and has lost the immortalizing oncogene [76]. Cath.a cells secrete dopamine and norepinephrine [86], and cells secreting hormones and neurotransmitters are specialized for secreting EVs [32]. Conversely, the N2a-PK1 cell line is derived from neuroblastoma Neuro 2A cells that produce low levels of tyrosine hydroxylase and dopamine unless they are differentiated [87]. It could be possible that the release of EVs is also specific to the cell type. The RK13 lines, which express PrP^C^ from sheep, mice, or voles infected by different prion strains, release PrP^TSE^ in exosomes differently [88].

## Conclusion

Prion protein is enriched in both large and small EVs. CAD5-RML lEVs, pelleted by 20,000× g and enriched in β-1 integrin, contain higher prion converting activity and infect native CAD5 cells more strongly than sEVs, pelleted by 110,000× g and depleted in β-1 integrin. We confirmed these results using three different methods and two approaches to infection, one assessing the concentration of infectivity in isolated EVs, and the second reflecting the total amount of infectivity associated with the EV fractions. For validation, we used the N2a-PK1-RML cell line, in which we also achieved stronger infection by lEVs. If the higher prion converting activity of lEVs was the sole reason why the infection with lEVs was stronger remains to be elucidated since there is limited knowledge of the differences upon uptake of different fractions of EVs. Small EVs or exosomes have been extensively studied compared to microvesicles, yet microvesicles could play significant roles in other diseases, such as cancer or cellular communication and immunity. Our data suggest the importance of lEVs in the trafficking of prions, which should be considered in future prion transmission studies.

## Declarations

**Availability of data and material:** We have submitted all relevant data of our experiments to the EV-TRACK knowledgebase (EV-TRACK ID: EV220318). The original data are available at the BioImage Archive repository (URL: https://www.ebi.ac.uk/biostudies/studies/S-BSST1019).

**The authors declare no conflicts of interest.**

**Funding:** The study was funded by Charles University projects GA UK 360216 and Cooperatio 207032-3 and The project National Institute for Neurological Research (Program EXCELES, ID Project No. LX22NPO5107) - Funded by the European Union – Next Generation EU.

**Author Contributions:** Conceptualization: Jakub Soukup, Methodology: Jakub Soukup, Cryo-electron Microscopy: Sami Kereïche, Jakub Soukup, RT-QuIC: Tibor Moško, writing – original draft: Jakub Soukup, writing – review and editing: Karel Holada, Sami Kereïche, Jakub Soukup, Funding: Jakub Soukup, Karel Holada.

### Acknowledgment

The authors thank Charles Weissmann from The Scripps Research Institute (FL, USA) for providing the CAD5, Peter Klöhn from MRC Prion Unit (London, UK) for providing the N2a-PK1 cells, the TSE Resource Centre (Roslin Institute, UK) for providing the AH6 antibody, and Adriano Aguzzi (Institute of Neuropathology, University of Zurich, Switzerland) for kindly providing the RML5 scrapie strain. The TEM was performed in the Vinicna Microscopy Core Facility co-financed by the Czech-BioImaging large RI project LM2023050. Computational resources were supplied by the project “e-Infrastruktura CZ” (e-INFRA LM2018140) provided within the program Projects of Large Research, Development and Innovations Infrastructure.

## References

[1] D.W. Colby, S.B. Prusiner, Prions, Cold Spring Harbor Perspectives in Biology 3 (2011) a006833. 10.1101/cshperspect.a006833.

[2] V.A. Lawson, S.J. Collins, C.L. Masters, A.F. Hill, Prion protein glycosylation, J. Neurochem. 93 (2005) 793-801. 10.1111/j.1471-4159.2005.03104.x.

[3] M. Kostelanska, J. Freisleben, Z.B. Hanusova, T. Mosko, R. Vik, D. Moravcova, A. Hamacek, J. Mosinger, K. Holada, Optimization of the photodynamic inactivation of prions by a phthalocyanine photosensitizer: The crucial involvement of singlet oxygen, J. Biophotonics 12 (2019) 13 e201800340. 10.1002/jbio.201800430.

[4] N. Stahl, D.R. Borchelt, K. Hsiao, S.B. Prusiner, SCRAPIE PRION PROTEIN CONTAINS A PHOSPHATIDYLINOSITOL GLYCOLIPID, Cell 51 (1987) 229–240. 10.1016/0092-8674(87)90150-4.

[5] R. Riek, G. Wider, M. Billeter, S. Hornemann, R. Glockshuber, K. Wuthrich, Prion protein NMR structure and familial human spongiform encephalopathies, Proceedings of the National Academy of Sciences of the United States of America 95 (1998) 11667–11672. 10.1073/pnas.95.20.11667.

[6] R. Zahn, A.Z. Liu, T. Luhrs, R. Riek, C. von Schroetter, F.L. Garcia, M. Billeter, L. Calzolai, G. Wider, K. Wuthrich, NMR solution structure of the human prion protein, Proceedings of the National Academy of Sciences of the United States of America 97 (2000) 145–150. 10.1073/pnas.97.1.145.

[7] M.P. McKinley, D.C. Bolton, S.B. Prusiner, A protease-resistant protein is a structural component of the Scrapie prion, Cell 35 (1983) 57–62. https://doi.org/10.1016/0092-8674(83)90207-6.

[8] K.C. Gough, B.C. Maddison, Prion transmission Prion excretion and occurrence in the environment, Prion 4 (2010) 275–282. 10.4161/pri.4.4.13678.

[9] P.A. McBride, W.J. Schulz-Schaeffer, M. Donaldson, M. Bruce, H. Diringer, H.A. Kretzschmar, M. Beekes, Early spread of scrapie from the gastrointestinal tract to the central nervous system involves autonomic fibers of the splanchnic and vagus nerves, Journal of Virology 75 (2001) 9320–9327. 10.1128/jvi.75.19.9320-9327.2001.

[10] B.R. Glaysher, N.A. Mabbott, Role of the GALT in scrapie agent neuroinvasion from the intestine, Journal of Immunology 178 (2007) 3757–3766. 10.4049/jimmunol.178.6.3757.

[11] K.L. Brown, K. Stewart, D.L. Ritchie, N.A. Mabbott, A. Williams, H. Fraser, W.I. Morrison, M.E. Bruce, Scrapie replication in lymphoid tissues depends on prion protein-expressing follicular dendritic cells, Nature Medicine 5 (1999) 1308–1312. 10.1038/15264.

[12] N. Kanu, Y. Imokawa, D.N. Drechsel, R.A. Williamson, C.R. Birkett, C.J. Bostock, J.P. Brockes, Transfer of scrapie prion infectivity by cell contact in culture, Curr. Biol. 12 (2002) 523–530 Pii s0960-9822(02)00722-4. 10.1016/s0960-9822(02)00722-4.

[13] K. Gousset, E. Schiff, C. Langevin, Z. Marijanovic, A. Caputo, D.T. Browman, N. Chenouard, F. de Chaumont, A. Martino, J. Enninga, J.C. Olivo-Marin, D. Mannel, C. Zurzolo, Prions hijack tunnelling nanotubes for intercellular spread, Nature Cell Biology 11 (2009) 328–U232. 10.1038/ncb1841.

[14] B. Fevrier, D. Vilette, F. Archer, D. Loew, W. Faigle, M. Vidal, H. Laude, G. Raposo, Cells release prions in association with exosomes, Proceedings of the National Academy of Sciences of the United States of America 101 (2004) 9683–9688. 10.1073/pnas.0308413101.

[15] G. Natale, M. Ferrucci, G. Lazzeri, A. Paparelli, F. Fornai, Transmission of prions within the gut and toward the central nervous system, Prion 5 (2011) 142–149. 10.4161/pri.5.3.16328.

[16] C.J. Sigurdson, J.C. Bartz, M. Glatzel, Cellular and Molecular Mechanisms of Prion Disease, Annu Rev Pathol 14 (2019) 497–516. 10.1146/annurev-pathmechdis-012418-013109.

[17] G. van Niel, D.R.F. Carter, A. Clayton, D.W. Lambert, G. Raposo, P. Vader, Challenges and directions in studying cell-cell communication by extracellular vesicles, Nat. Rev. Mol. Cell Biol. 23 (2022) 369–382. 10.1038/s41580-022-00460-3.

[18] M. Colombo, G. Raposo, C. Thery, Biogenesis, Secretion, and Intercellular Interactions of Exosomes and Other Extracellular Vesicles, in: R. Schekman, R. Lehmann (Eds.) Annual Review of Cell and Developmental Biology, Vol 30, Annual Reviews, Palo Alto, 2014, pp. 255-289.

[19] W.M. Henne, N.J. Buchkovich, S.D. Emr, The ESCRT Pathway, Dev. Cell 21 (2011) 77-91. 10.1016/j.devcel.2011.05.015.

[20] K. Trajkovic, C. Hsu, S. Chiantia, L. Rajendran, D. Wenzel, F. Wieland, P. Schwille, B. Brugger, M. Simons, Ceramide triggers budding of exosome vesicles into multivesicular Endosomes, Science 319 (2008) 1244–1247. 10.1126/science.1153124.

[21] H.F.G. Heijnen, A.E. Schiel, R. Fijnheer, H.J. Geuze, J.J. Sixma, Activated platelets release two types of membrane vesicles: Microvesicles by surface shedding and exosomes derived from exocytosis of multivesicular bodies and alpha-granules, Blood 94 (1999) 3791–3799. 10.1182/blood.V94.11.3791.423a22_3791_3799.

[22] S.I. Buschow, E.N.M. Nolte-’t Hoen, G. van Niel, M.S. Pols, T. ten Broeke, M. Lauwen, F. Ossendorp, C.J.M. Melief, G. Raposo, R. Wubbolts, M.H.M. Wauben, W. Stoorvogel, MHC II in Dendritic Cells is Targeted to Lysosomes or T Cell-Induced Exosomes Via Distinct Multivesicular Body Pathways, Traffic 10 (2009) 1528–1542. 10.1111/j.1600-0854.2009.00963.x.

[23] J. Klumperman, G. Raposo, The Complex Ultrastructure of the Endolysosomal System, Cold Spring Harbor Perspectives in Biology 6 (2014) 22 a016857. 10.1101/cshperspect.a016857.

[24] N.P. Hessvik, A. verbye, A. Brech, M.L. Torgersen, I.S. Jakobsen, K. Sandvig, A. Llorente, PIKfyve inhibition increases exosome release and induces secretory autophagy, Cellular and Molecular Life Sciences 73 (2016) 4717–4737. 10.1007/s00018-016-2309-8.

[25] L. Murrow, R. Malhotra, J. Debnath, ATG12-ATG3 interacts with Alix to promote basal autophagic flux and late endosome function, Nature Cell Biology 17 (2015) 300-+. 10.1038/ncb3112.

[26] M.F. Baietti, Z. Zhang, E. Mortier, A. Melchior, G. Degeest, A. Geeraerts, Y. Ivarsson, F. Depoortere, C. Coomans, E. Vermeiren, P. Zimmermann, G. David, Syndecan-syntenin-ALIX regulates the biogenesis of exosomes, Nature Cell Biology 14 (2012) 677–685. 10.1038/ncb2502.

[27] J. Kowal, G. Arras, M. Colombo, M. Jouve, J.P. Morath, B. Primdal-Bengtson, F. Dingli, D. Loew, M. Tkach, C. Thery, Proteomic comparison defines novel markers to characterize heterogeneous populations of extracellular vesicle subtypes, Proceedings of the National Academy of Sciences of the United States of America 113 (2016) E968–E977. 10.1073/pnas.1521230113.

[28] J. Wang, X.J. Zhuang, K.S. Greene, H. Si, M.A. Antonyak, J.E. Druso, K.F. Wilson, R.A. Cerione, Q.Y. Feng, H.Y. Wang, Cdc42 functions as a regulatory node for tumour-derived microvesicle biogenesis, Journal of Extracellular Vesicles 10 (2021) 17 e12051. 10.1002/jev2.12051.

[29] H.J. Dai, S.S. Zhang, X.K. Du, W.K. Zhang, R. Jing, X.X. Wang, L.H. Pan, RhoA inhibitor suppresses the production of microvesicles and rescues high ventilation induced lung injury, Int. Immunopharmacol. 72 (2019) 74–81. 10.1016/j.intimp.2019.03.059.

[30] M.J. Sun, X.F. Xue, L.Y. Li, D.D. Xu, S.H. Li, S.W.C. Li, Q.N. Su, Ectosome biogenesis and release processes observed by using live-cell dynamic imaging in mammalian glial cells, Quant. Imaging Med. Surg. 11 (2021) 4604–4616. 10.21037/qims-20-1015.

[31] J.F. Nabhan, R.X. Hu, R.S. Oh, S.N. Cohen, Q. Lu, Formation and release of arrestin domain-containing protein 1-mediated microvesicles (ARMMs) at plasma membrane by recruitment of TSG101 protein, Proceedings of the National Academy of Sciences of the United States of America 109 (2012) 4146–4151. 10.1073/pnas.1200448109.

[32] G. van Niel, G. D’Angelo, G. Raposo, Shedding light on the cell biology of extracellular vesicles, Nat. Rev. Mol. Cell Biol. 19 (2018) 213–228. 10.1038/nrm.2017.125.

[33] Z. Andreu, M. Yanez-Mo, Tetraspanins in extracellular vesicle formation and function, Front. Immunol. 5 (2014) 12 442. 10.3389/fimmu.2014.00442.

[34] V. Laghezza Masci, A.R. Taddei, G. Gambellini, F. Giorgi, A.M. Fausto, Microvesicles shed from fibroblasts act as metalloproteinase carriers in a 3-D collagen matrix, J Circ Biomark 5 (2016) 1849454416663660. 10.1177/1849454416663660.

[35] F. Momen-Heravi, S.J. Getting, S.A. Moschos, Extracellular vesicles and their nucleic acids for biomarker discovery, Pharmacol. Ther. 192 (2018) 170–187. 10.1016/j.pharmthera.2018.08.002.

[36] D.K. Jeppesen, A.M. Fenix, J.L. Franklin, J.N. Higginbotham, Q. Zhang, L.J. Zimmerman, D.C. Liebler, J. Ping, Q. Liu, R. Evans, W.H. Fissell, J.G. Patton, L.H. Rome, D.T. Burnette, R.J. Coffey, Reassessment of Exosome Composition, Cell 177 (2019) 428-+. 10.1016/j.cell.2019.02.029.

[37] A.M. Deleo, T. Ikezu, Extracellular Vesicle Biology in Alzheimer’s Disease and Related Tauopathy, J. Neuroimmune Pharm. 13 (2018) 292–308. 10.1007/s11481-017-9768-z.

[38] A. Bobrie, M. Colombo, S. Krumeich, G.A. Raposo, C. Thery, Diverse subpopulations of vesicles secreted by different intracellular mechanisms are present in exosome preparations obtained by differential ultracentrifugation, Journal of Extracellular Vesicles 1 (2012) 11 18397. 10.3402/jev.v1i0.18397.

[39] H.Y. Zhang, D. Freitas, H.S. Kim, K. Fabijanic, Z. Li, H.Y. Chen, M.T. Mark, H. Molina, A.B. Martin, L. Bojmar, J. Fang, S. Rampersaud, A. Hoshino, I. Matei, C.M. Kenific, M. Nakajima, A.P. Mutvei, P. Sansone, W. Buehring, H.J. Wang, J.P. Jimenez, L. Cohen-Gould, N. Paknejad, M. Brendel, K. Manova-Todorova, A. Magalhaes, J.A. Ferreira, H. Osorio, A.M. Silva, A. Massey, J.R. Cubillos-Ruiz, G. Galletti, P. Giannakakou, A.M. Cuervo, J. Blenis, R. Schwartz, M.S. Brady, H. Peinado, J. Bromberg, H. Matsui, C.A. Reis, D. Lyden, Identification of distinct nanoparticles and subsets of extracellular vesicles by asymmetric flow field-flow fractionation, Nature Cell Biology 20 (2018) 332-+. 10.1038/s41556-018-0040-4.

[40] C. Sunyach, A. Jen, J. Deng, K.T. Fitzgerald, Y. Frobert, J. Grassi, M.W. McCaffrey, R. Morris, The mechanism of internalization of glycosylphosphatidylinositol-anchored prion protein, Embo Journal 22 (2003) 3591–3601. 10.1093/emboj/cdg344.

[41] D. Sarnataro, V. Campana, S. Paladino, M. Stornaiuolo, L. Nitsch, C. Zurzolo, PrPC association with lipid rafts in the early secretory pathway stabilizes its cellular conformation, Molecular Biology of the Cell 15 (2004) 4031–4042. 10.1091/mbc.E03-05-0271.

[42] Z. Fremuntova, T. Mosko, J. Soukup, J. Kucerova, M. Kostelanska, Z.B. Hanusova, M. Filipova, L. Cervenakova, K. Holada, Changes in cellular prion protein expression, processing and localisation during differentiation of the neuronal cell line CAD 5, Biology of the Cell 112 (2020) 1–21. 10.1111/boc.201900045.

[43] A. Brouckova, K. Holada, Cellular prion protein in blood platelets associates with both lipid rafts and the cytoskeleton, Thromb. Haemost. 102 (2009) 966–974. 10.1160/th09-02-0074.

[44] A. Didonna, Prion protein and its role in signal transduction, Cell. Mol. Biol. Lett. 18 (2013) 209–230. 10.2478/s11658-013-0085-0.

[45] S.L. Shyng, M.T. Huber, D.A. Harris, A PRION PROTEIN CYCLES BETWEEN THE CELL-SURFACE AND AN ENDOCYTIC COMPARTMENT IN CULTURED NEUROBLASTOMA-CELLS, Journal of Biological Chemistry 268 (1993) 15922–15928.

[46] P.J. Peters, A. Mironov, D. Peretz, E. van Donselaar, E. Leclerc, S. Erpel, S.J. DeArmond, D.R. Burton, R.A. Williamson, M. Vey, S.B. Prusiner, Trafficking of prion proteins through a caveolae-mediated endosomal pathway, J. Cell Biol. 162 (2003) 703–717. 10.1083/jcb.200304140.

[47] Y.I. Yim, B.C. Park, R. Yadavalli, X.H. Zhao, E. Eisenberg, L.E. Greene, The multivesicular body is the major internal site of prion conversion, J. Cell Sci. 128 (2015) 1434–1443. 10.1242/jcs.165472.

[48] B.B. Guo, S.A. Bellingham, A.F. Hill, The Neutral Sphingomyelinase Pathway Regulates Packaging of the Prion Protein into Exosomes, Journal of Biological Chemistry 290 (2015) 3455–3467. 10.1074/jbc.M114.605253.

[49] N.M. Veith, H. Plattner, C.A.O. Stuermer, W.J. Schulz-Schaeffer, A. Burkle, Immunolocalisation of PrPSc in scrapie-infected N2a mouse neuroblastoma cells by light and electron microscopy, Eur. J. Cell Biol. 88 (2009) 45–63. 10.1016/j.ejcb.2008.08.001.

[50] Z. Marijanovic, A. Caputo, V. Campana, C. Zurzolo, Identification of an Intracellular Site of Prion Conversion, Plos Pathogens 5 (2009) 15 e1000426. 10.1371/journal.ppat.1000426.

[51] D. Vilette, K. Laulagnier, A. Huor, S. Alais, S. Simoes, R. Maryse, M. Provansal, S. Lehmann, O. Andreoletti, L. Schaeffer, G. Raposo, P. Leblanc, Efficient inhibition of infectious prions multiplication and release by targeting the exosomal pathway, Cellular and Molecular Life Sciences 72 (2015) 4409–4427. 10.1007/s00018-015-1945-8.

[52] B.B. Guo, S.A. Bellingham, A.F. Hill, Stimulating the Release of Exosomes Increases the Intercellular Transfer of Prions, Journal of Biological Chemistry 291 (2016) 5128–5137. 10.1074/jbc.M115.684258.

[53] L.J. Vella, R.A. Sharples, V.A. Lawson, C.L. Masters, R. Cappai, A.F. Hill, Packaging of prions into exosomes is associated with a novel pathway of PrP processing, Journal of Pathology 211 (2007) 582–590. 10.1002/path.2145.

[54] G. D’Arrigo, M. Gabrielli, F. Scaroni, P. Swuec, L. Amin, A. Pegoraro, E. Adinolfi, F. Di Virgilio, D. Cojoc, G. Legname, C. Verderio, Astrocytes-derived extracellular vesicles in motion at the neuron surface: Involvement of the prion protein, Journal of Extracellular Vesicles 10 (2021) 22 e12114. 10.1002/jev2.12114.

[55] R. Goold, S. Rabbanian, L. Sutton, R. Andre, P. Arora, J. Moonga, A.R. Clarke, G. Schiavo, P. Jat, J. Collinge, S.J. Tabrizi, Rapid cell-surface prion protein conversion revealed using a novel cell system, Nat. Commun. 2 (2011) 281. 10.1038/ncomms1282.

[56] T. Yamasaki, G.S. Baron, A. Suzuki, R. Hasebe, M. Horiuchi, Characterization of intracellular dynamics of inoculated PrP-res and newly generated PrPSc during early stage prion infection in Neuro2a cells, Virology 450 (2014) 324–335. 10.1016/j.virol.2013.11.007.

[57] R. Goold, C. McKinnon, S. Rabbanian, J. Collinge, G. Schiavo, S.J. Tabrizi, Alternative fates of newly formed PrPSc upon prion conversion on the plasma membrane, J. Cell Sci. 126 (2013) 3552–3562. 10.1242/jcs.120477.

[58] V. Mattei, M.G. Barenco, V. Tasciotti, T. Garofalo, A. Longo, K. Boller, J. Loewer, R. Misasi, F. Montrasio, M. Sorice, Paracrine Diffusion of PrPC and Propagation of Prion Infectivity by Plasma Membrane-Derived Microvesicles, Plos One 4 (2009) e5057. 10.1371/journal.pone.0005057.

[59] P. Saa, O. Yakovleva, J. de Castro, I. Vasilyeva, S.H. De Paoli, J. Simak, L. Cervenakova, First Demonstration of Transmissible Spongiform Encephalopathy-associated Prion Protein (PrPTSE) in Extracellular Vesicles from Plasma of Mice Infected with Mouse-adapted Variant Creutzfeldt-Jakob Disease by in Vitro Amplification, Journal of Biological Chemistry 289 (2014) 29247–29260. 10.1074/jbc.M114.589564.

[60] L. Cervenakova, P. Saa, O. Yakoyleva, I. Vasilyeva, J. de Castro, P. Brown, R. Dodd, Are prions transported by plasma exosomes?, Transfusion and Apheresis Science 55 (2016) 70–83. 10.1016/j.transci.2016.07.013.

[61] P. Saa, Is sporadic Creutzfeldt-Jakob disease transfusion-transmissible?, Transfusion 60 (2020) 655–658. 10.1111/trf.15763.

[62] A. Benmoussa, C.H.C. Lee, B. Laffont, P. Savard, J. Laugier, E. Boilard, C. Gilbert, I. Fliss, P. Provost, Commercial Dairy Cow Milk microRNAs Resist Digestion under Simulated Gastrointestinal Tract Conditions, J. Nutr. 146 (2016) 2206–2215. 10.3945/jn.116.237651.

[63] M.M. Rahman, K. Shimizu, M. Yamauchi, H. Takase, S. Ugawa, A. Okada, Y. Inoshima, Acidification effects on isolation of extracellular vesicles from bovine milk, Plos One 14 (2019) 12 e0222613. 10.1371/journal.pone.0222613.

[64] C. Thery, K.W. Witwer, E. Aikawa, M.J. Alcaraz, J.D. Anderson, R. Andriantsitohaina, A. Antoniou, T. Arab, F. Archer, G.K. Atkin-Smith, D.C. Ayre, J.M. Bach, D. Bachurski, H. Baharvand, L. Balaj, S. Baldacchino, N.N. Bauer, A.A. Baxter, M. Bebawy, C. Beckham, A.B. Zavec, A. Benmoussa, A.C. Berardi, P. Bergese, E. Bielska, C. Blenkiron, S. Bobis-Wozowicz, E. Boilard, W. Boireau, A. Bongiovanni, F.E. Borras, S. Bosch, C.M. Boulanger, X. Breakefield, A.M. Breglio, M.A. Brennan, D.R. Brigstock, A. Brisson, M.L.D. Broekman, J.F. Bromberg, P. Bryl-Gorecka, S. Buch, A.H. Buck, D. Burger, S. Busatto, D. Buschmann, B. Bussolati, E.I. Buzas, J.B. Byrd, G. Camussi, D.R.F. Carter, S. Caruso, L.W. Chamley, Y.T. Chang, C.C. Chen, S. Chen, L. Cheng, A.R. Chin, A. Clayton, S.P. Clerici, A. Cocks, E. Cocucci, R.J. Coffey, A. Cordeiro-da-Silva, Y. Couch, F.A.W. Coumans, B. Coyle, R. Crescitelli, M.F. Criado, C. D’Souza-Schorey, S. Das, A.D. Chaudhuri, P. de Candia, E.F. De Santana, O. De Wever, H.A. del Portillo, T. Demaret, S. Deville, A. Devitt, B. Dhondt, D. Di Vizio, L.C. Dieterich, V. Dolo, A.P.D. Rubio, M. Dominici, M.R. Dourado, T.A.P. Driedonks, F.V. Duarte, H.M. Duncan, R.M. Eichenberger, K. Ekstrom, S.E.L. Andaloussi, C. Elie-Caille, U. Erdbrugger, J.M. Falcon-Perez, F. Fatima, J.E. Fish, M. Flores-Bellver, A. Forsonits, A. Frelet-Barrand, F. Fricke, G. Fuhrmann, S. Gabrielsson, A. Gamez-Valero, C. Gardiner, K. Gartner, R. Gaudin, Y.S. Gho, B. Giebel, C. Gilbert, M. Gimona, I. Giusti, D.C.I. Goberdhan, A. Gorgens, S.M. Gorski, D.W. Greening, J.C. Gross, A. Gualerzi, G.N. Gupta, D. Gustafson, A. Handberg, R.A. Haraszti, P. Harrison, H. Hegyesi, A. Hendrix, A.F. Hill, F.H. Hochberg, K.F. Hoffmann, B. Holder, H. Holthofer, B. Hosseinkhani, G.K. Hu, Y.Y. Huang, V. Huber, S. Hunt, A.G.E. Ibrahim, T. Ikezu, J.M. Inal, M. Isin, A. Ivanova, H.K. Jackson, S. Jacobsen, S.M. Jay, M. Jayachandran, G. Jenster, L.Z. Jiang, S.M. Johnson, J.C. Jones, A. Jong, T. Jovanovic-Talisman, S. Jung, R. Kalluri, S. Kano, S. Kaur, Y. Kawamura, E.T. Keller, D. Khamari, E. Khomyakova, A. Khvorova, P. Kierulf, K.P. Kim, T. Kislinger, M. Klingeborn, D.J. Klinke, M. Kornek, M.M. Kosanovic, A.F. Kovacs, E.M. Kramer-Albers, S. Krasemann, M. Krause, I.V. Kurochkin, G.D. Kusuma, S. Kuypers, S. Laitinen, S.M. Langevin, L.R. Languino, J. Lannigan, C. Lasser, L.C. Laurent, G. Lavieu, E. Lazaro-Ibanez, S. Le Lay, M.S. Lee, Y.X.F. Lee, D.S. Lemos, M. Lenassi, A. Leszczynska, I.T.S. Li, K. Liao, S.F. Libregts, E. Ligeti, R. Lim, S.K. Lim, A. Line, K. Linnemannstons, A. Llorente, C.A. Lombard, M.J. Lorenowicz, A.M. Lorincz, J. Lotvall, J. Lovett, M.C. Lowry, X. Loyer, Q. Lu, B. Lukomska, T.R. Lunavat, S.L.N. Maas, H. Malhi, A. Marcilla, J. Mariani, J. Mariscal, E.S. Martens-Uzunova, L. Martin-Jaular, M.C. Martinez, V.R. Martins, M. Mathieu, S. Mathivanan, M. Maugeri, L.K. McGinnis, M.J. McVey, D.G. Meckes, K.L. Meehan, I. Mertens, V.R. Minciacchi, A. Moller, M.M. Jorgensen, A. Morales-Kastresana, J. Morhayim, F. Mullier, M. Muraca, L. Musante, V. Mussack, D.C. Muth, K.H. Myburgh, T. Najrana, M. Nawaz, I. Nazarenko, P. Nejsum, C. Neri, T. Neri, R. Nieuwland, L. Nimrichter, J.P. Nolan, E.N.M. Nolte-’t Hoen, N. Noren Hooten, L. O’Driscoll, T. O’Grady, A. O’Loghlen, T. Ochiya, M. Olivier, A. Ortiz, L.A. Ortiz, X. Osteikoetxea, O. Ostegaard, M. Ostrowski, J. Park, D.M. Pegtel, H. Peinado, F. Perut, M.W. Pfaffl, D.G. Phinney, B.C.H. Pieters, R.C. Pink, D.S. Pisetsky, E.P. von Strandmann, I. Polakovicova, I.K.H. Poon, B.H. Powell, I. Prada, L. Pulliam, P. Quesenberry, A. Radeghieri, R.L. Raffai, S. Raimondo, J. Rak, M.I. Ramirez, G. Raposo, M.S. Rayyan, N. Regev-Rudzki, F.L. Ricklefs, P.D. Robbins, D.D. Roberts, S.C. Rodrigues, E. Rohde, S. Rome, K.M.A. Rouschop, A. Rughetti, A.E. Russell, P. Saa, S. Sahoo, E. Salas-Huenuleo, C. Sanchez, J.A. Saugstad, M.J. Saul, R.M. Schiffelers, R. Schneider, T.H. Schoyen, A. Scott, E. Shahaj, S. Sharma, O. Shatnyeva, F. Shekari, G.V. Shelke, A.K. Shetty, K. Shiba, P.R.M. Siljander, A.M. Silva, A. Skowronek, O.L. Snyder, R.P. Soares, B.W. Sodar, C. Soekmadji, J. Sotillo, P.D. Stahl, W. Stoorvogel, S.L. Stott, E.F. Strasser, S. Swift, H. Tahara, M. Tewari, K. Timms, S. Tiwari, R. Tixeira, M. Tkach, W.S. Toh, R. Tomasini, A.C. Torrecilhas, J.P. Tosar, V. Toxavidis, L. Urbanelli, P. Vader, B.W.M. van Balkom, S.G. van der Grein, J. Van Deun, M.J.C. van Herwijnen, K. Van Keuren-Jensen, G. van Niel, M.E. van Royen, A.J. van Wijnen, M.H. Vasconcelos, I.J. Vechetti, T.D. Veit, L.J. Vella, E. Velot, F.J. Verweij, B. Vestad, J.L. Vinas, T. Visnovitz, K.V. Vukman, J. Wahlgren, D.C. Watson, M.H.M. Wauben, A. Weaver, J.P. Webber, V. Weber, A.M. Wehman, D.J. Weiss, J.A. Welsh, S. Wendt, A.M. Wheelock, Z. Wiener, L. Witte, J. Wolfram, A. Xagorari, P. Xander, J. Xu, X.M. Yan, M. Yanez-Mo, H. Yin, Y. Yuana, V. Zappulli, J. Zarubova, V. Zekas, J.Y. Zhang, Z.Z. Zhao, L. Zheng, A.R. Zheutlin, A.M. Zickler, P. Zimmermann, A.M. Zivkovic, D. Zocco, E.K. Zuba-Surma, Minimal information for studies of extracellular vesicles 2018 (MISEV2018): a position statement of the International Society for Extracellular Vesicles and update of the MISEV2014 guidelines, Journal of Extracellular Vesicles 7 (2018) 43 1535750. 10.1080/20013078.2018.1535750.

[65] S.P. Mahal, C.A. Baker, C.A. Demczyk, E.W. Smith, C. Julius, C. Weissmann, Prion strain discrimination in cell culture: The cell panel assay, Proceedings of the National Academy of Sciences 104 (2007) 20908–20913. doi:10.1073/pnas.0710054104.

[66] P.C. Klohn, L. Stoltze, E. Flechsig, M. Enari, C. Weissmann, A quantitative, highly sensitive cell-based infectivity assay for mouse scrapie prions, Proceedings of the National Academy of Sciences of the United States of America 100 (2003) 11666–11671. 10.1073/pnas.1834432100.

[67] J. Van Deun, P. Mestdagh, P. Agostinis, Ö. Akay, S. Anand, J. Anckaert, Z.A. Martinez, T. Baetens, E. Beghein, L. Bertier, G. Berx, J. Boere, S. Boukouris, M. Bremer, D. Buschmann, J.B. Byrd, C. Casert, L. Cheng, A. Cmoch, D. Daveloose, E. De Smedt, S. Demirsoy, V. Depoorter, B. Dhondt, T.A.P. Driedonks, A. Dudek, A. Elsharawy, I. Floris, A.D. Foers, K. Gärtner, A.D. Garg, E. Geeurickx, J. Gettemans, F. Ghazavi, B. Giebel, T.G. Kormelink, G. Hancock, H. Helsmoortel, A.F. Hill, V. Hyenne, H. Kalra, D. Kim, J. Kowal, S. Kraemer, P. Leidinger, C. Leonelli, Y. Liang, L. Lippens, S. Liu, A. Lo Cicero, S. Martin, S. Mathivanan, P. Mathiyalagan, T. Matusek, G. Milani, M. Monguió-Tortajada, L.M. Mus, D.C. Muth, A. Németh, E.N.M. Nolte-’t Hoen, L. O’Driscoll, R. Palmulli, M.W. Pfaffl, B. Primdal-Bengtson, E. Romano, Q. Rousseau, S. Sahoo, N. Sampaio, M. Samuel, B. Scicluna, B. Soen, A. Steels, J.V. Swinnen, M. Takatalo, S. Thaminy, C. Théry, J. Tulkens, I. Van Audenhove, S. van der Grein, A. Van Goethem, M.J. van Herwijnen, G. Van Niel, N. Van Roy, A.R. Van Vliet, N. Vandamme, S. Vanhauwaert, G. Vergauwen, F. Verweij, A. Wallaert, M. Wauben, K.W. Witwer, M.I. Zonneveld, O. De Wever, J. Vandesompele, A. Hendrix, E.-T. Consortium, EV-TRACK: transparent reporting and centralizing knowledge in extracellular vesicle research, Nature Methods 14 (2017) 228–232. 10.1038/nmeth.4185.

[68] C. Théry, S. Amigorena, G. Raposo, A. Clayton, Isolation and Characterization of Exosomes from Cell Culture Supernatants and Biological Fluids, Current Protocols in Cell Biology 30 (2006) 3.22.21-23.22.29. https://doi.org/10.1002/0471143030.cb0322s30.

[69] O. Janouskova, J. Rakusan, M. Karaskova, K. Holada, Photodynamic inactivation of prions by disulfonated hydroxyaluminium phthalocyanine, Journal of General Virology 93 (2012) 2512–2517. 10.1099/vir.0.044727-0.

[70] A. Foutz, B.S. Appleby, C. Hamlin, X.Q. Liu, S. Yang, Y. Cohen, W. Chen, J. Blevins, C. Fausett, H. Wang, P. Gambetti, S.L. Zhang, A. Hughson, C. Tatsuoka, L.B. Schonberger, M.L. Cohen, B. Caughey, J.G. Safar, Diagnostic and prognostic value of human prion detection in cerebrospinal fluid, Ann. Neurol. 81 (2017) 79–92. 10.1002/ana.24833.

[71] T. Mosko, S. Galuskova, R. Matej, M. Buzova, K. Holada, Detection of Prions in Brain Homogenates and CSF Samples Using a Second-Generation RT-QuIC Assay: A Useful Tool for Retrospective Analysis of Archived Samples, Pathogens 10 (2021) 13 750. 10.3390/pathogens10060750.

[72] Z. Hanusova, T. Mosko, R. Matej, K. Holada, Precision in the design of an experimental study deflects the significance of proteinase-activated receptor 2 expression in scrapie-inoculated mice, Journal of General Virology 98 (2017) 1563–1569. 10.1099/jgv.0.000803.

[73] J. Soukup, M. Kostelanska, S. Kereiche, A. Hujacova, M. Pavelcova, J. Petrak, E.K. Havrdova, K. Holada, Flow Cytometry Analysis of Blood Large Extracellular Vesicles in Patients with Multiple Sclerosis Experiencing Relapse of the Disease, J. Clin. Med. 11 (2022) 15 2832. 10.3390/jcm11102832.

[74] J. Dubochet, M. Adrian, J.J. Chang, J.C. Homo, J. Lepault, A.W. McDowall, P. Schultz, Cryo-electron microscopy of vitrified specimens, Q Rev Biophys 21 (1988) 129–228. 10.1017/s0033583500004297.

[75] S.F. Godsave, H. Wille, J. Pierson, S.B. Prusiner, P.J. Peters, Plasma membrane invaginations containing clusters of full-length PrPSc are an early form of prion-associated neuropathology in vivo, Neurobiol. Aging 34 (2013) 1621–1631. 10.1016/j.neurobiolaging.2012.12.015.

[76] Y.P. Qi, J.K.T. Wang, M. McMillian, D.M. Chikaraishi, Characterization of a CNS cell line, CAD, in which morphological differentiation is initiated by serum deprivation, J. Neurosci. 17 (1997) 1217–1225.

[77] P.D. Stahl, G. Raposo, Extracellular Vesicles: Exosomes and Microvesicles, Integrators of Homeostasis, Physiology 34 (2019) 169-177. 10.1152/physiol.00045.2018.

[78] S. Gurung, D. Perocheau, L. Touramanidou, J. Baruteau, The exosome journey: from biogenesis to uptake and intracellular signalling, Cell Communication and Signaling 19 (2021) 47. 10.1186/s12964-021-00730-1.

[79] M. Mathieu, N. Nevo, M. Jouve, J.I. Valenzuela, M. Maurin, F.J. Verweij, R. Palmulli, D. Lankar, F. Dingli, D. Loew, E. Rubinstein, G. Boncompain, F. Perez, C. Thery, Specificities of exosome versus small ectosome secretion revealed by live intracellular tracking of CD63 and CD9, Nat. Commun. 12 (2021) 18 4389. 10.1038/s41467-021-24384-2.

[80] T. Takizawa, C. Tatematsu, K. Watanabe, K. Kato, Y. Nakanishi, Cleavage of calnexin caused by apoptotic stimuli: Implication for the regulation of apoptosis, J. Biochem.(Tokyo) 136 (2004) 399–405. 10.1093/jb/mvh133.

[81] X. Osteikoetxea, A. Balogh, K. Szabó-Taylor, A. Németh, T.G. Szabó, K. Pálóczi, B. Sódar, Á. Kittel, B. György, É. Pállinger, J. Matkó, E.I. Buzás, Improved Characterization of EV Preparations Based on Protein to Lipid Ratio and Lipid Properties, PLOS ONE 10 (2015) e0121184. 10.1371/journal.pone.0121184.

[82] J. de Vrij, S.L. Maas, M. van Nispen, M. Sena-Esteves, R.W. Limpens, A.J. Koster, S. Leenstra, M.L. Lamfers, M.L. Broekman, Quantification of nanosized extracellular membrane vesicles with scanning ion occlusion sensing, Nanomedicine (Lond) 8 (2013) 1443–1458. 10.2217/nnm.12.173.

[83] F.A.W. Coumans, E. van der Pol, A.N. Böing, N. Hajji, G. Sturk, T.G. van Leeuwen, R. Nieuwland, Reproducible extracellular vesicle size and concentration determination with tunable resistive pulse sensing, Journal of Extracellular Vesicles 3 (2014) 25922. 10.3402/jev.v3.25922.

[84] E. van der Pol, F.A. Coumans, A.E. Grootemaat, C. Gardiner, I.L. Sargent, P. Harrison, A. Sturk, T.G. van Leeuwen, R. Nieuwland, Particle size distribution of exosomes and microvesicles determined by transmission electron microscopy, flow cytometry, nanoparticle tracking analysis, and resistive pulse sensing, J Thromb Haemost 12 (2014) 1182–1192. 10.1111/jth.12602.

[85] R. Morales, P.P. Hu, C. Duran-Aniotz, F. Moda, R. Diaz-Espinoza, B. Chen, J. Bravo-Alegria, N. Makarava, I.V. Baskakov, C. Soto, Strain-dependent profile of misfolded prion protein aggregates, Sci Rep 6 (2016) 20526. 10.1038/srep20526.

[86] C. Suri, B.P. Fung, A.S. Tischler, D.M. Chikaraishi, Catecholaminergic cell lines from the brain and adrenal glands of tyrosine hydroxylase-SV40 T antigen transgenic mice, J Neurosci 13 (1993) 1280–1291. 10.1523/jneurosci.13-03-01280.1993.

[87] R.G. Tremblay, M. Sikorska, J.K. Sandhu, P. Lanthier, M. Ribecco-Lutkiewicz, M. Bani-Yaghoub, Differentiation of mouse Neuro 2A cells into dopamine neurons, J Neurosci Methods 186 (2010) 60–67. 10.1016/j.jneumeth.2009.11.004.

[88] Z.E. Arellano-Anaya, A. Huor, P. Leblanc, S. Lehmann, M. Provansal, G. Raposo, O. Andreoletti, D. Vilette, Prion strains are differentially released through the exosomal pathway, Cellular and Molecular Life Sciences 72 (2015) 1185–1196. 10.1007/s00018-014-1735-8.

