## Supplementary figures and tables for "Large Extracellular Vesicles Transfer Higher Levels of Prions and Infect Cell Culture Better than Small Extracellular Vesicles"

### 1 Supplementary Figures and Tables

#### 1.1 Supplementary Figures

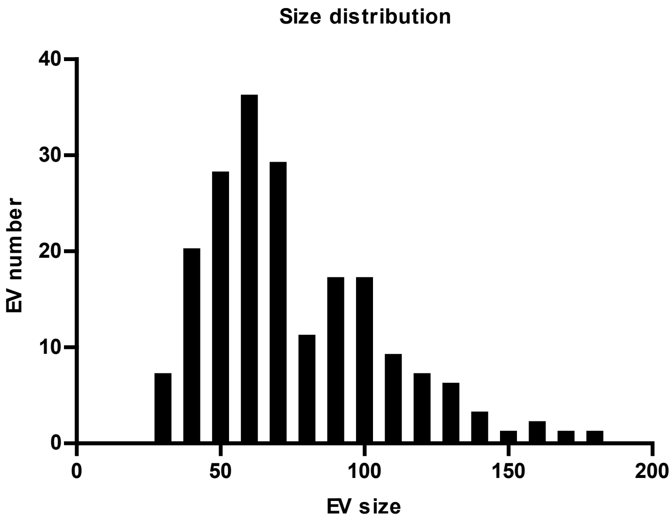


**Supplementary Figure S1: Size distribution of EVs present in sEV fraction.** Sizes were measured in ImageJ software on cryo-EM images.

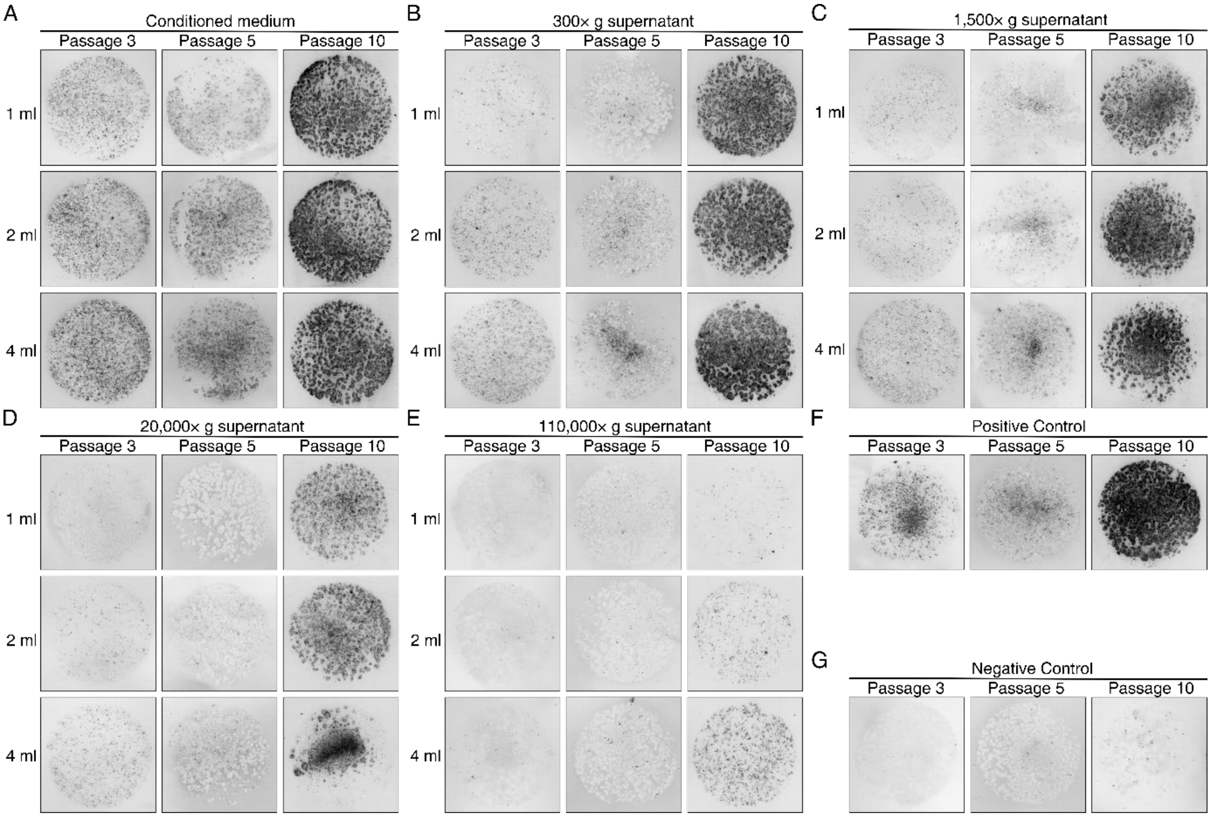


**Supplementary Figure S2: Representative cell blots from infection of CAD5 cells with supernatants collected during isolation steps of EVs.** (a) Infection with whole conditioned medium. (b) Infection with supernatant after pelleting of dead cells (300× g). (c) Infection with supernatant after pelleting cell debris and apoptotic bodies (1,500× g). (d) Infection with supernatant after pelleting lEVs (20,000 × g). (e) Infection with supernatant after pelleting sEVs (110,000× g). (f) Positive control made from CAD5-RML cell homogenate. (g) Negative control made from native CAD5 cell homogenate. For visualization purposes, all images were enhanced with brightness and contrast.

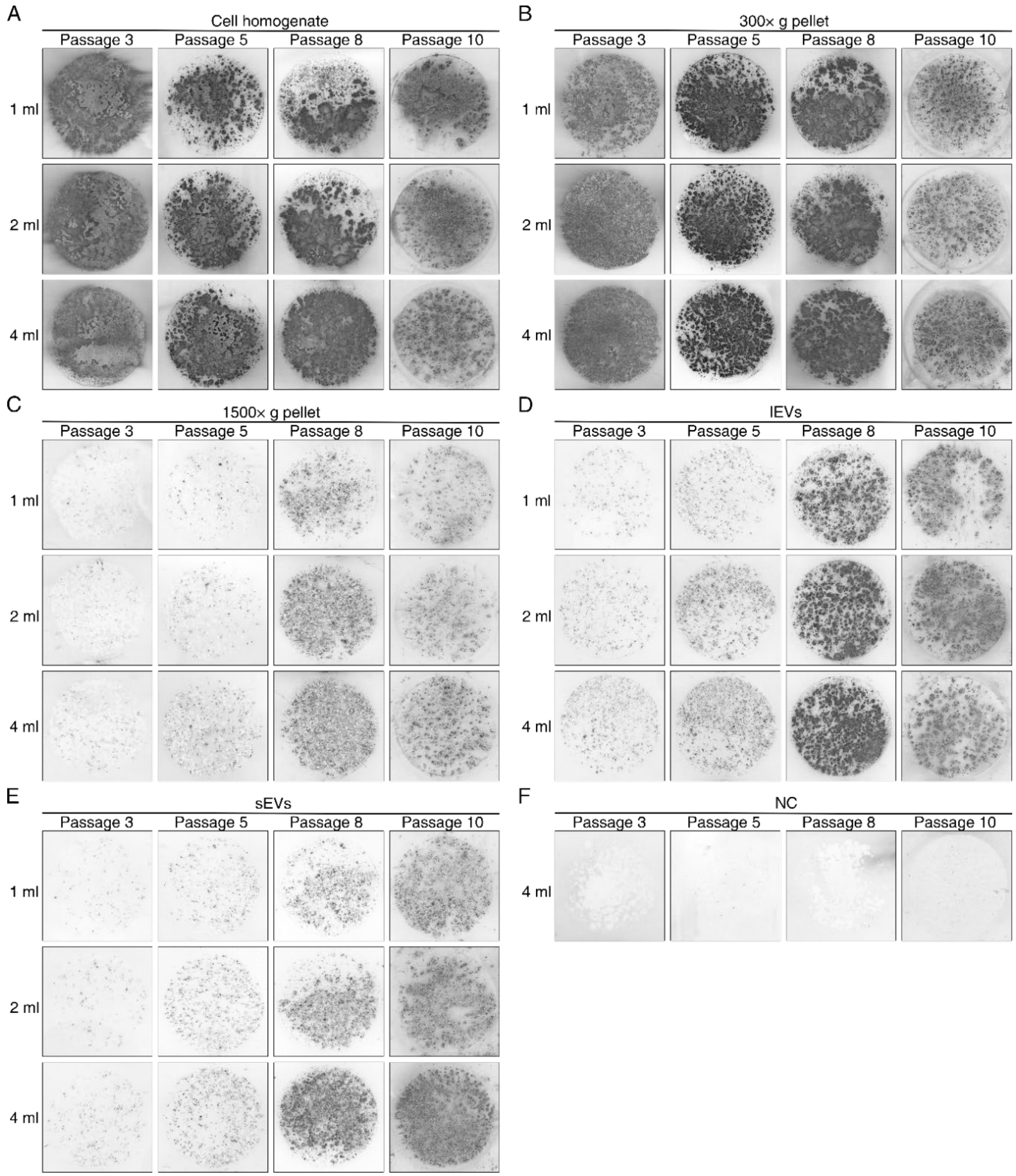


**Supplementary Figure S3: Representative cell blots from the infection of CAD5 cells with pellets obtained during EV isolation steps.** The infection was performed according to the OVS scheme, and 4, 2, or 1 ml of suspension was used for infection. (d, e, and f are part of Fig. 4 and are shown here for comparison with other fractions. (a) Cells harvested after collection of conditioned medium, homogenized, and resuspended in fresh medium. (b) Pellet representing dead cells in conditioning medium (300× g). (c) Pellet representing large debris and apoptotic bodies (1,500× g). (d) Pelleted lEVs (20,000× g). (e) Pelleted sEVs (110,000× g). (f) Negative control. Harvested and homogenized native CAD5 cells. For visualization purposes, all images were enhanced with brightness and contrast.

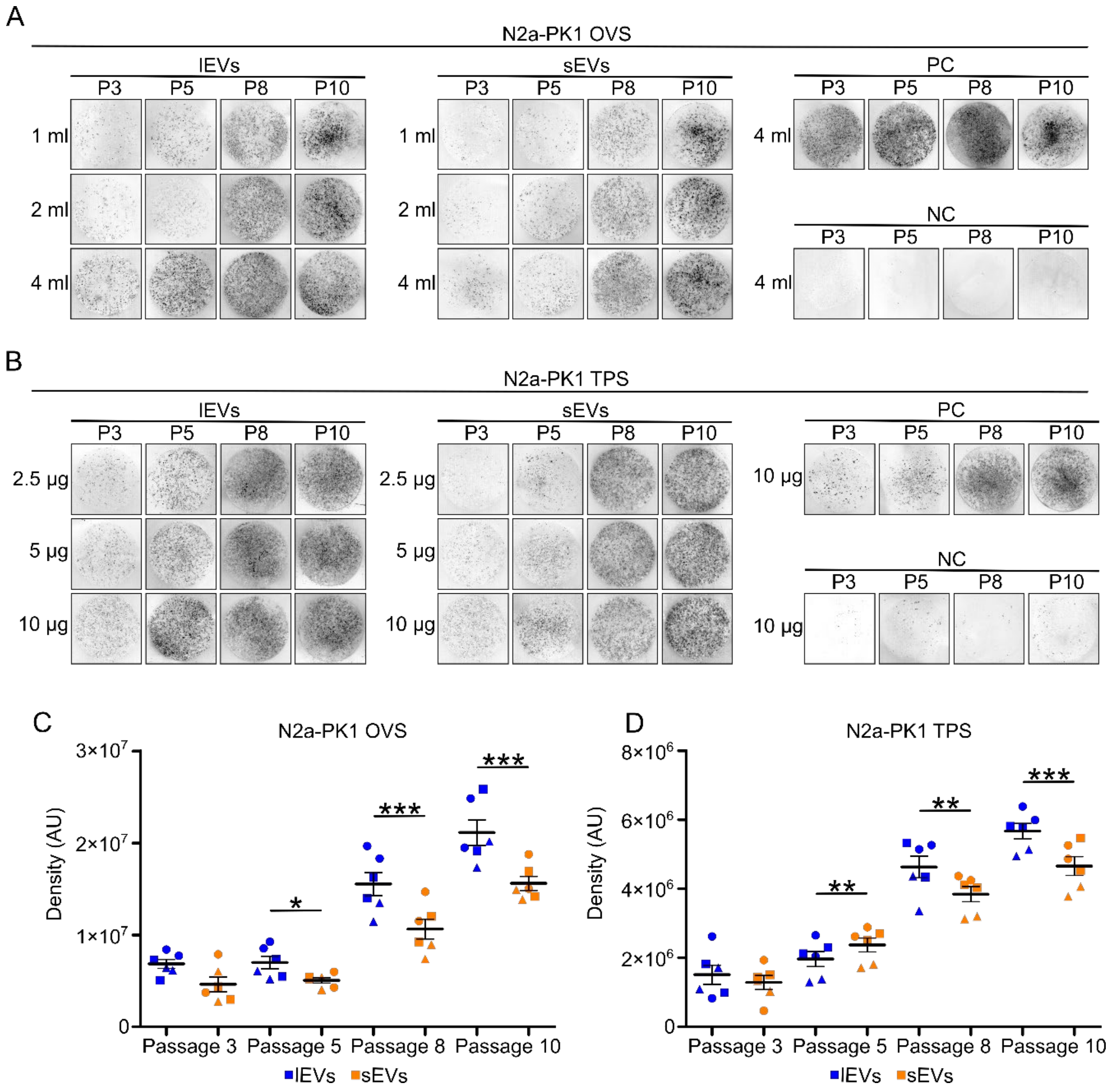


**Supplementary Figure S4: Example images of cell blots from passages after infection of N2a-PK1 cells with lEVs and sEVs with densitometry analysis.** (a and b) Example images of cell blots from infection of native N2a-PK1 cells infected with lEVs and sEVs in different passages after infection (passage 3, 5, 8, and 10). Cells were infected with 4 ml, 2 ml, and 1 ml of resuspended EVs in OVS (a) infection or with 10 µg, 5 µg, and 2.5 µg of EVs in TPS infection (b). NC – negative control, PC – positive control. For visualization purposes, all images were enhanced with brightness and contrast. (c and d) Densitometry analysis of cell blots from different passages. Samples were analyzed by two-tailed non-parametric paired t-test (Wilcoxon matched-pairs signed rank test) with pairing of corresponding infection dose. The line represents the mean with SEM. Blue spots – lEV infected cells, orange spots – sEV infected cells, circles – infection with 4 ml or 10 µg, squares - infection with 2 ml or 5 µg, triangles - infection with 1 ml or 2.5 µg. Each spot is average of technical duplicate. * $P < 0.05$; ** $P < 0.01$, *** $P < 0.001$. AU – arbitrary units.

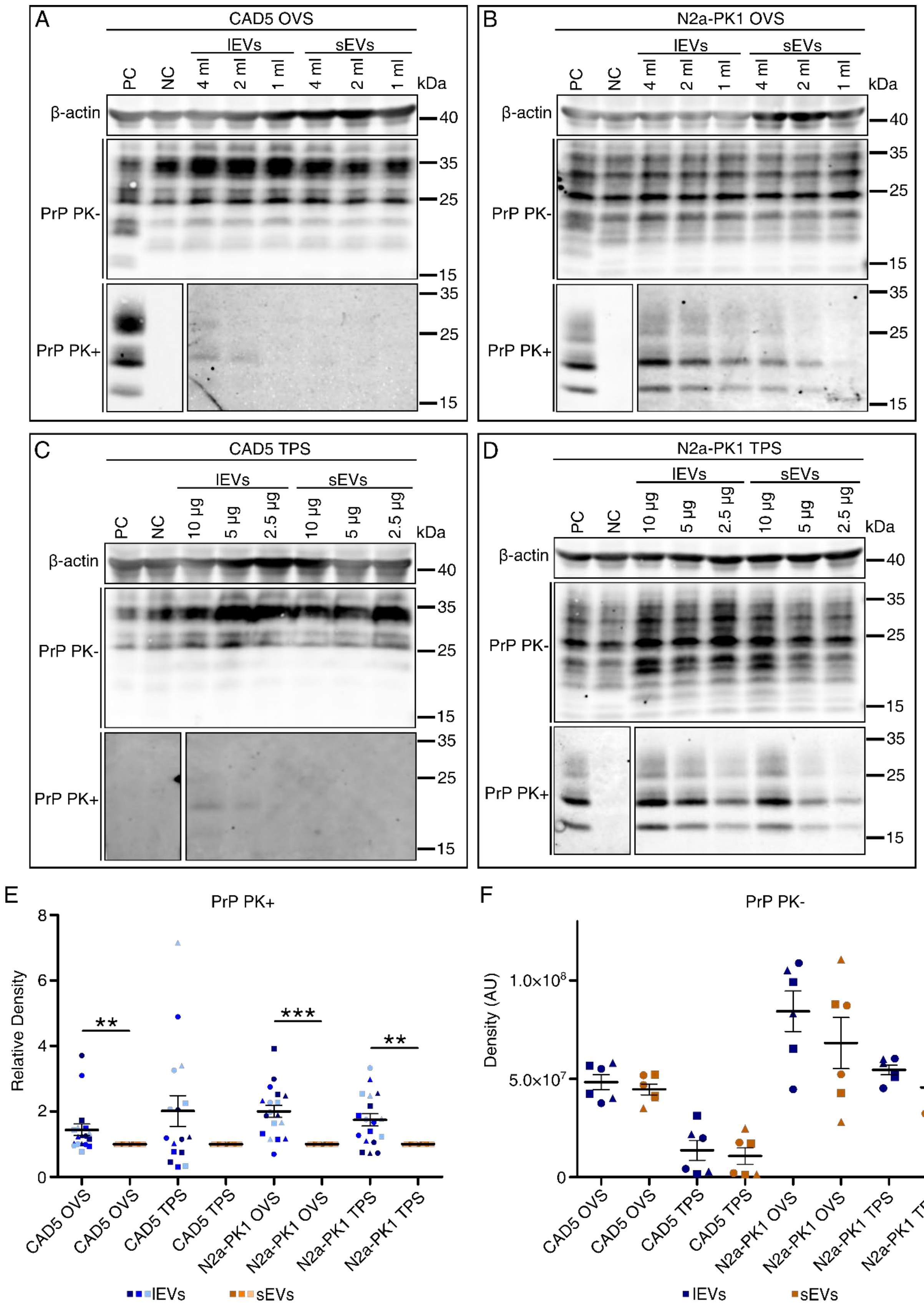

A
CAD5 OVS
lEVs
sEVs
PC
NC
4 ml
2 ml
1 ml
kDa
β-actin
PrP PK-
PrP PK+
40
35
25
15
B
N2a-PK1 OVS
C
CAD5 TPS
10 µg
5 µg
2.5 µg
D
N2a-PK1 TPS
E
PrP PK+
Relative Density
CAD5 OVS
CAD5 TPS
N2a-PK1 OVS
N2a-PK1 TPS
**
***
**
lEVs
sEVs
F
PrP PK-
Density (AU)
1.0×10^8
5.0×10^7
0

**Supplementary Figure S5: western blot analysis of PrPres and $PrP^{Sc}$/$PrP^{C}$signals in cells infected by lEV and sEV fractions at passage 3.** (a-d) An example images of western blots from infection of CAD5 (a, c) or N2a-PK1 (b, d) cells with lEVs and sEVs at passage 3 after infection. Cells were infected with 4 ml, 2 ml, and 1 ml of resuspended EVs in OVS infection (a, b) or 10 µg, 5 µg, and 2.5 µg of washed EVs in TPS infection (c, d). PK+ - proteinase K digested samples. PK- - proteinase K untreated samples. β-Actin was used as a loading control for PrP PK- samples, which were also compared. All samples are shown for a loading of 40 µg per line. NC – negative control (mock-infected cells), PC – positive control. For visualization purposes, all images were enhanced with brightness and contrast. (e) Relative comparison of PrP signal in PrP PK+ samples by densitometry. The line represents the mean with SEM. Blue colors – lEVs, orange colors – sEVs. Light blue/orange – loading 10 µg per line, blue/orange – loading 20 µg per line, dark blue/orange – loading 40 µg per line. Circles – infection with 4 ml or 10 µg, squares - infection with 2 ml or 5 µg, triangles - infection with 1 ml or 2.5 µg. Samples were analyzed by two-tailed non-parametric paired t-test (Wilcoxon matched-pairs signed rank test) with the pairing of corresponding infection dose and protein loading per line. * $P < 0.05$; ** $P < 0.01$, *** $P < 0.001$. The quantification precision is limited in the CAD5 TPS infection – the PrPres signal of infection by sEVs is not visible yet. (f) Comparison of PrP PK- bands density. Blue spots – lEV infected cells, orange spots – sEV infected cells, circles – infection dose 4 ml or 10 µg, squares - infection dose 2 ml or 5 µg, triangles - infection dose 4 ml or 10 µg. The same statistical approach was used for PrP PK- analysis. AU – arbitrary units.

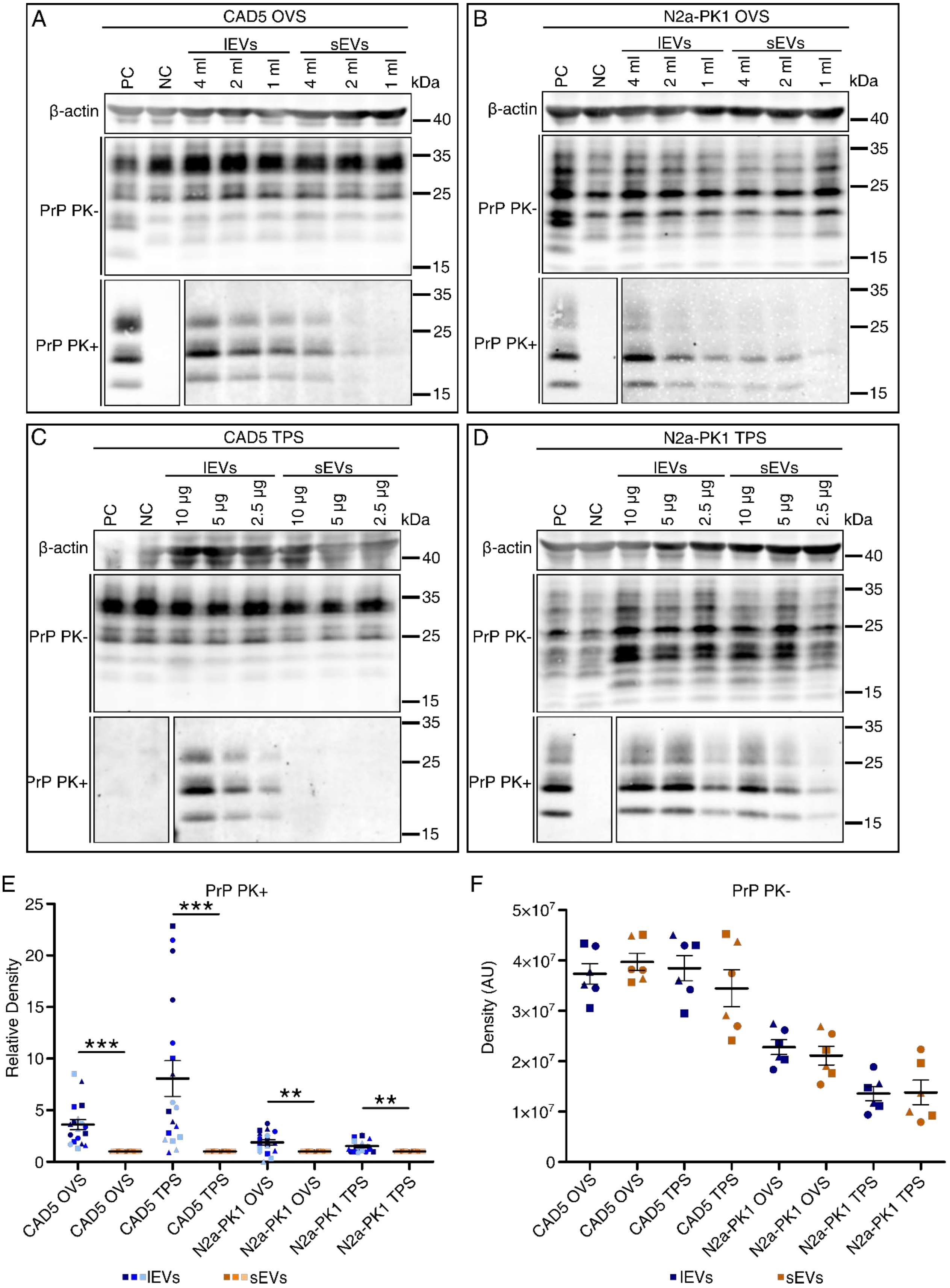
A
CAD5 OVS
lEVs
sEVs
PC
NC
4 ml
2 ml
1 ml
4 ml
2 ml
1 ml
kDa
β-actin
PrP PK-
PrP PK+
40
35
25
15
B
N2a-PK1 OVS
C
CAD5 TPS
10 µg
5 µg
2.5 µg
D
N2a-PK1 TPS
E
PrP PK+
Relative Density
***
**
CAD5 OVS
CAD5 TPS
N2a-PK1 OVS
N2a-PK1 TPS
lEVs
sEVs
F
PrP PK-
Density (AU)
5×10^7
4×10^7
3×10^7
2×10^7
1×10^7

**Supplementary Figure S6: western blot analysis of PrPres and $PrP^{Sc}/PrP^{C}$signals in cells infected by lEV and sEV fractions at passage 5.** (a-d) An example images of western blots from infection of CAD5 (a, c) or N2a-PK1 (b, d) cells with lEVs and sEVs at passage 5 after infection. Cells were infected with 4 ml, 2 ml, and 1 ml of resuspended EVs in OVS infection (a, b) or 10 µg, 5 µg, and 2.5 µg of washed EVs in TPS infection (c, d). PK+ - proteinase K digested samples. PK- - proteinase K untreated samples. β-Actin was used as a loading control for PrP PK- samples, which were also compared. All samples are shown for a loading of 40 µg per line. NC – negative control (mock-infected cells), PC – positive control. For visualization purposes, all images were enhanced with brightness and contrast. (e) Relative comparison of PrP signal in PrP PK+ samples by densitometry. The line represents the mean with SEM. Blue colors – lEVs, orange colors – sEVs. Light blue/orange – loading 10 µg per line, blue/orange – loading 20 µg per line, dark blue/orange – loading 40 µg per line. Circles – infection with 4 ml or 10 µg, squares - infection with 2 ml or 5 µg, triangles - infection with 1 ml or 2.5 µg. Samples were analyzed by two-tailed non-parametric paired t-test (Wilcoxon matched-pairs signed rank test) with the pairing of corresponding infection dose and protein loading per line. * $P < 0.05$; ** $P < 0.01$, *** $P < 0.001$. The quantification precision is limited in the CAD5 TPS infection – the PrPres signal of infection by sEVs is not visible yet. (f) Comparison of PrP PK- bands density. Blue spots – lEV infected cells, orange spots – sEV infected cells, circles – infection dose 4 ml or 10 µg, squares - infection dose 2 ml or 5 µg, triangles - infection dose 4 ml or 10 µg. The same statistical approach was used for PrP PK- analysis. AU – arbitrary units.

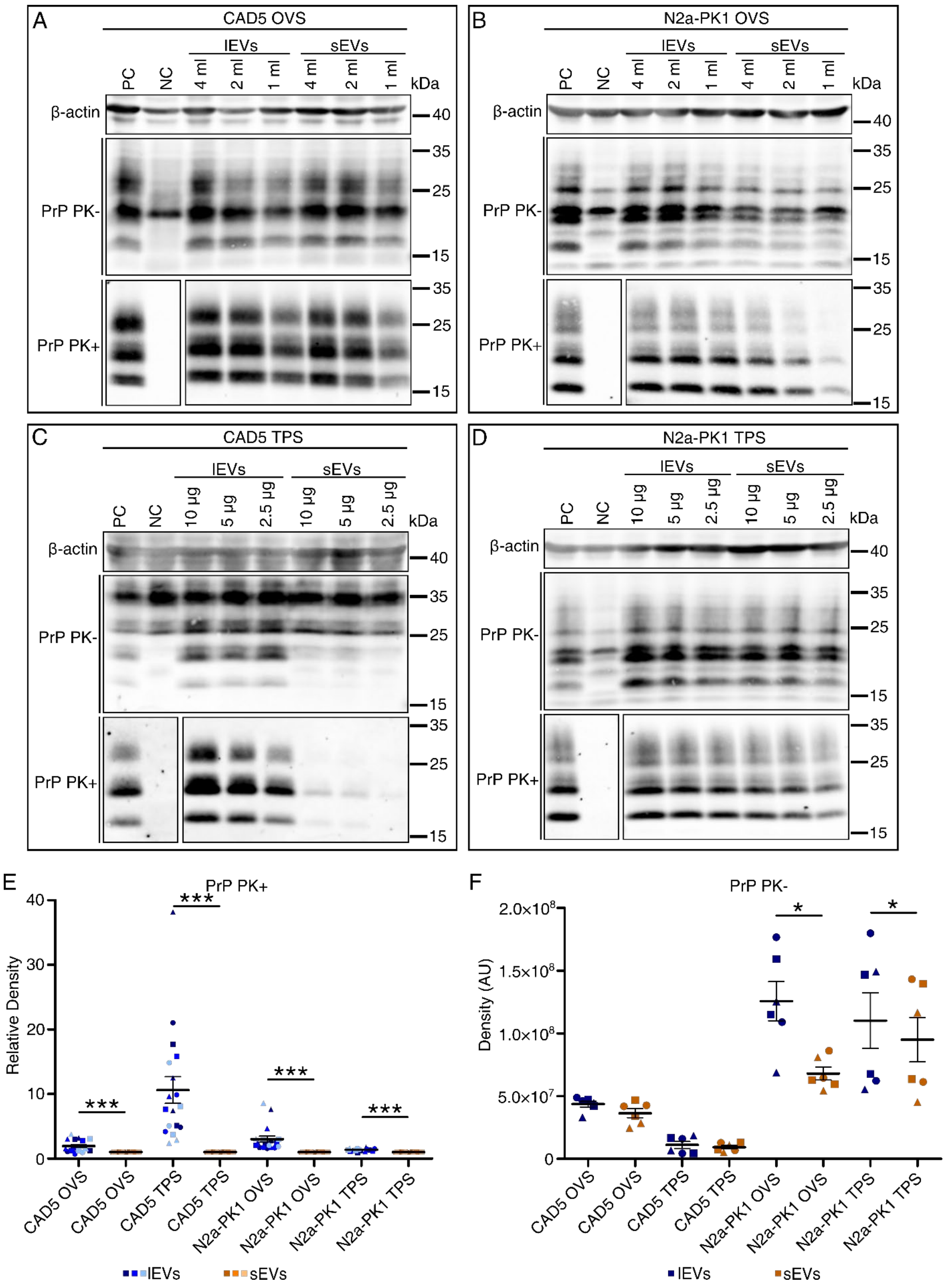
A
CAD5 OVS
lEVs
sEVs
PC
NC
4 ml
2 ml
1 ml
kDa
β-actin
PrP PK-
PrP PK+
B
N2a-PK1 OVS
C
CAD5 TPS
10 µg
5 µg
2.5 µg
D
N2a-PK1 TPS
E
PrP PK+
Relative Density
CAD5 OVS
CAD5 TPS
N2a-PK1 OVS
N2a-PK1 TPS
lEVs
sEVs
F
PrP PK-
Density (AU)
2.0x10^8
1.5x10^8
1.0x10^8
5.0x10^7

**Supplementary Figure S7: western blot analysis of PrPres and $PrP^{Sc}$/$PrP^{C}$signals in cells infected by lEV and sEV fractions at passage 10.** (a-d) An example images of western blots from infection of CAD5 (a, c) or N2a-PK1 (b, d) cells with lEVs and sEVs at passage 10 after infection. Cells were infected with 4 ml, 2 ml, and 1 ml of resuspended EVs in OVS infection (a, b) or 10 µg, 5 µg, and 2.5 µg of washed EVs in TPS infection (c, d). PK+ - proteinase K digested samples. PK- - proteinase K untreated samples. β-Actin was used as a loading control for PrP PK- samples, which were also compared. All samples are shown for a loading of 40 µg per line. NC – negative control (mock-infected cells), PC – positive control. For visualization purposes, all images were enhanced with brightness and contrast. (e) Relative comparison of PrP signal in PrP PK+ samples by densitometry. The line represents the mean with SEM. Blue colors – lEVs, orange colors – sEVs. Light blue/orange – loading 10 µg per line, blue/orange – loading 20 µg per line, dark blue/orange – loading 40 µg per line. Circles – infection with 4 ml or 10 µg, squares - infection with 2 ml or 5 µg, triangles - infection with 1 ml or 2.5 µg. Samples were analyzed by two-tailed non-parametric paired t-test (Wilcoxon matched-pairs signed rank test) with the pairing of corresponding infection dose and protein loading per line. * $P < 0.05$; ** $P < 0.01$, *** $P < 0.001$. The quantification precision is limited in the CAD5 TPS infection – the PrPres in infection by lEVs was reaching the maximum. (f) Comparison of PrP PK- bands density. Blue spots – lEV infected cells, orange spots – sEV infected cells, circles – infection dose 4 ml or 10 µg, squares - infection dose 2 ml or 5 µg, triangles - infection dose 4 ml or 10 µg. The same statistical approach was used for PrP PK- analysis. AU – arbitrary units.

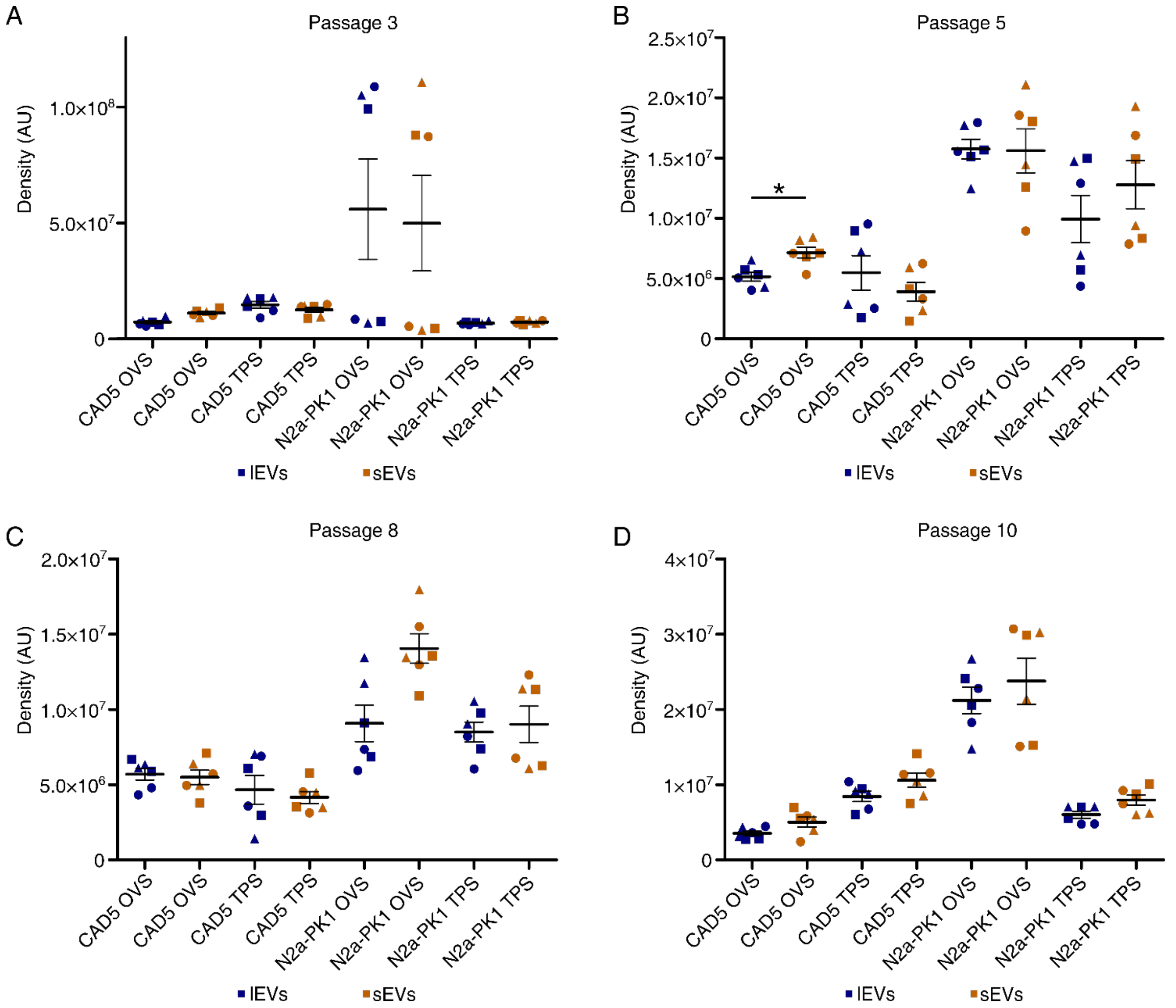


**Supplementary Figure S8: western blot analysis of β-actin loading control for Figure 6 and Supplementary Figures 5 - 7.** Each spot represents absolute density in arbitrary units (AU) of each β-actin band. Blue spots – lEV infected cells, orange spots – sEV infected cells, circles – infection dose 4 ml or 10 µg, squares - infection dose 2 ml or 5 µg, triangles - infection dose 4 ml or 10 µg. (a) passage 3 (control for Figure S5), (b) passage 5 (control for Figure S6), (c) passage 8 (control for Figure 6), (d) passage 10 (control for Figure S7). The line represents the mean with SEM. Samples were analyzed by two-tailed non-parametric paired t-test (Wilcoxon matched-pairs signed rank test) with pairing of corresponding infection dose. * $P < 0.05$.

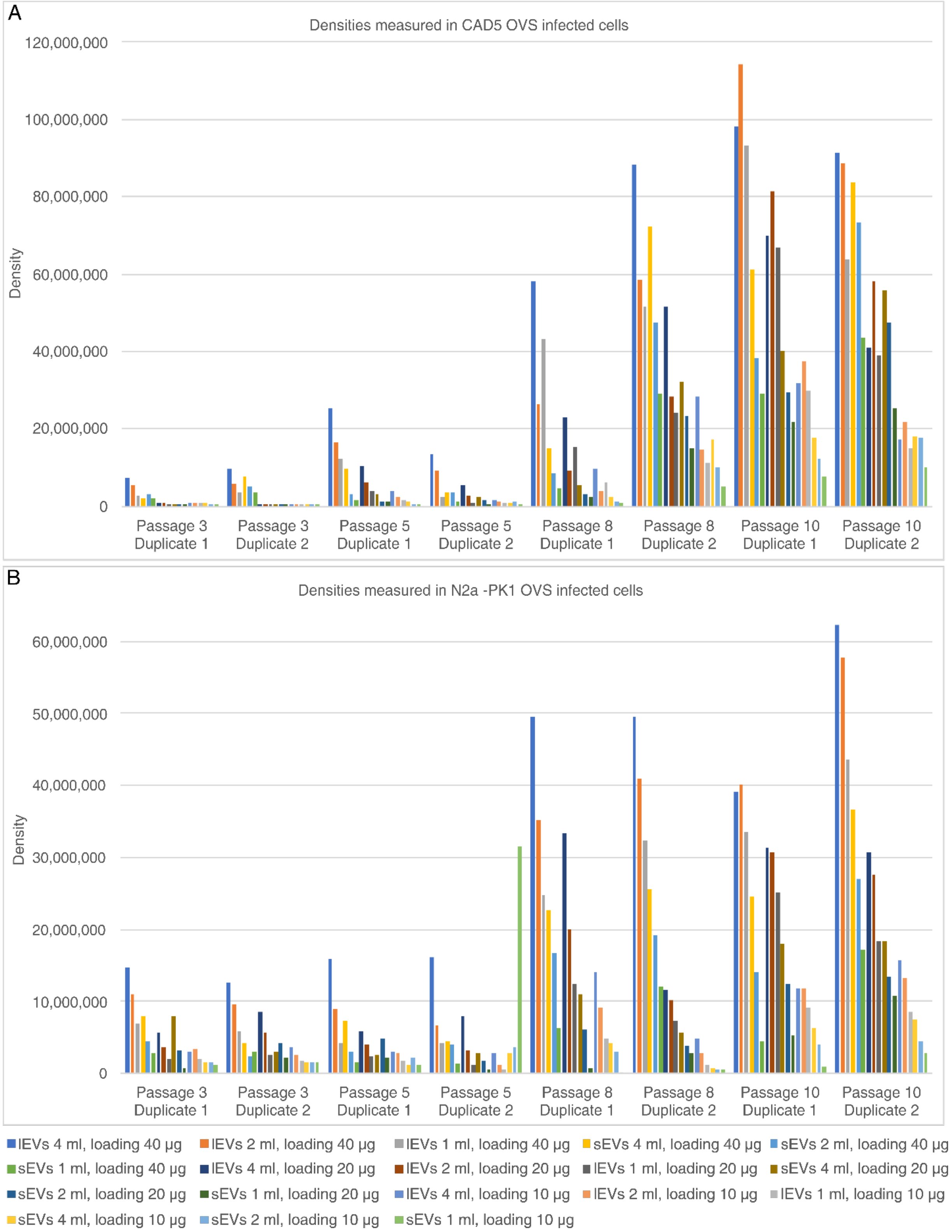


**Supplementary Figure S9: Absolute PrPres densities of western blots of OVS-infected cells.** (a, b) Densities measured in passage 3, 5, 8, and 10 after infection of CAD5 (a) or N2a-PK1 (b) cells with EV fractions. The cells were infected with 4 ml, 2 ml, and 1 ml of EVs resuspended in the original volume of the medium. Duplicates 1 and 2 represent two independent experiments.

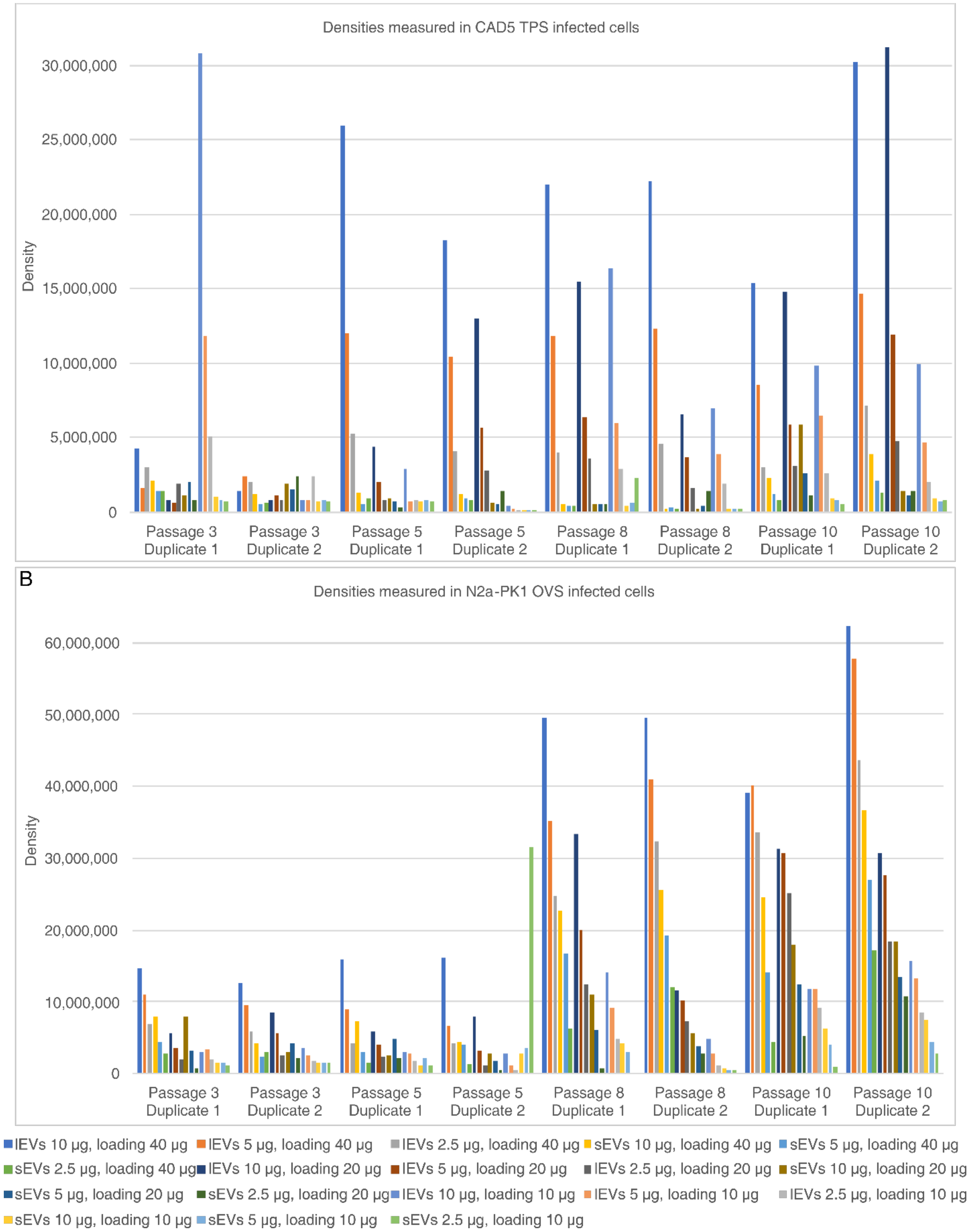


**Supplementary Figure S10: Absolute PrPres densities of western blots of TPS-infected cells.** (a, b) Densities measured in passage 3, 5, 8, and 10 after infection of CAD5 (a) or N2a-PK1 (b) cells with EV fractions. The cells were infected with 10 µg, 5 µg, and 2.5 µg of washed. Duplicates 1 and 2 represent two independent experiments.

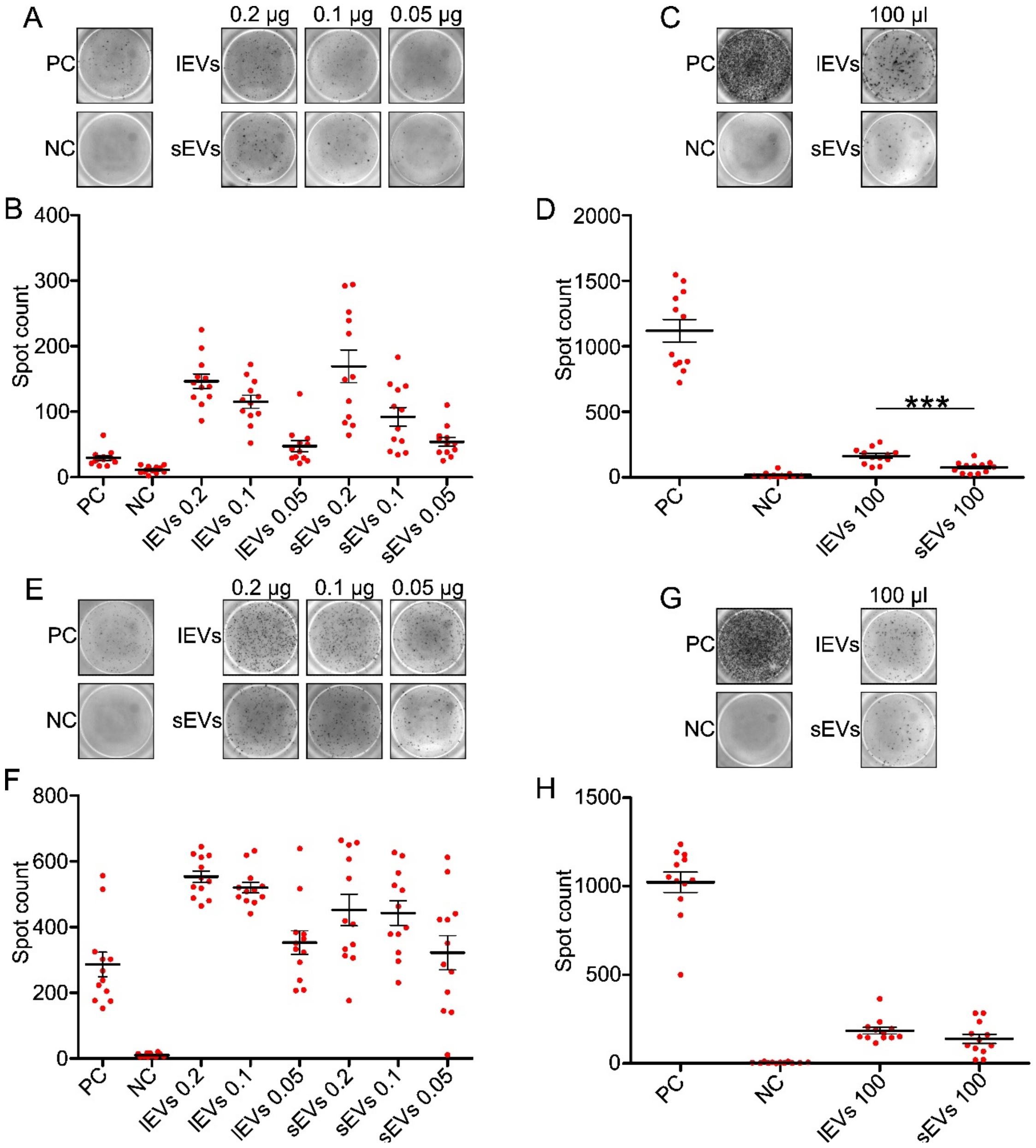


**Supplementary Figure S11: Scrapie cell assay of CAD5 cells infected with EVs.** (a, e) Example images of wells in the scrapie cell assay infected with the TPS scheme in passages 3 (a), and 8 (e). PC – positive control (infection by CAD5-RML cell homogenate), NC – negative control (infection by CAD5 cell homogenate). A total of 1000 cells were transferred to each well. (c, g) Representative images of wells in the scrapie cell assay infected with the OVS scheme in passages 3 (c), and 5 (g). A total of 10,000 cells were transferred to each well. For visualization purposes, all images were enhanced with brightness and contrast. (b, f) Spot count in each well of the scrapie cell assay (hexaplicates of infection from a duplicate of isolation) – TPS infection in passage 3 (b), and 8 (f). Each circle represents the count in one well. (d, h) Spot count in each well of the scrapie cell assay in OVS infection at passage 3 (d), and 5 (h). The thick line represents the mean value with SEM, statistical significance was determined using a two-tailed unpaired t-test. ***P < 0.001

### 1.2 Supplementary Tables

| Marker | Sample | CAD5-RML | N2a-PK1-RML |
|---|---|---|---|
| β-1 Integrin | Cell lysate | 18,297,712 | 29,860,517 |
| | lEVs | 4,867,254 | 12,491,569 |
| | sEVs | 960,115 | 4,571,642 |
| Calnexin | Cell lysate | 24,440,445 | 20,841,566 |
| | lEVs | 2,377,631 | 740,284 |
| | sEVs | 1,124,943 | 35,404 |
| Alix | Cell lysate | 181,258 | 119,094 |
| | lEVs | 2,810,074 | 29,300 |
| | sEVs | 562,786 | 125,868 |
| TSG-101 | Cell lysate | 237,535 | 1,192,251 |
| | lEVs | 193,699 | 382,167 |
| | sEVs | 3,044,397 | 5,632,658 |
| HSP70 | Cell lysate | 9,688,849 | 18,435,051 |
| | lEVs | 7,792,009 | 19,732,372 |
| | sEVs | 3,371,024 | 26,249,701 |
| CD63 | Cell lysate | 1,426,221 | 3,015,685 |
| | lEVs | 457,465 | 947,706 |
| | sEVs | 806,082 | 1,765,069 |
| CD9 | Cell lysate | 205,854 | 57,741 |
| | lEVs | 4,185,503 | 259,967 |
| | sEVs | 48,368,381 | 5,580,693 |

**Supplementary Table S1: Measured density in markers detected in Figure 1C and 1D.** The density was measured on 10 µg loading per line since it has better signal to background ratio.

CAD5-RML 1st isolation

| Sample | Dilution | Positive wells |
|---|---|---|
| cell lysate | $10^{-6}$ | 4/4 |
| | $10^{-7}$ | 4/4 |
| | $10^{-8}$ | 2/4 |
| | $10^{-9}$ | 0/4 |
| lEVs | $10^{-6}$ | 4/4 |
| | $10^{-7}$ | 4/4 |
| | $10^{-8}$ | 2/4 |
| | $10^{-9}$ | 0/4 |
| sEVs | $10^{-6}$ | 4/4 |
| | $10^{-7}$ | 1/4 |
| | $10^{-8}$ | 0/4 |
| | $10^{-9}$ | 0/4 |
| cell lysate (negative) | $10^{-6}$ | 0/4 |

CAD5-RML 2nd isolation

| Sample | Dilution | Positive wells |
|---|---|---|
| cell lysate | $10^{-5}$ | 4/4 |
| | $2\times10^{-6}$ | 4/4 |
| lEVs | $10^{-5}$ | 4/4 |
| | $2\times10^{-6}$ | 4/4 |
| | $4\times10^{-7}$ | 4/4 |
| | $8\times10^{-8}$ | 4/4 |
| | $2\times10^{-8}$ | 2/4 |
| | $3\times10^{-9}$ | 1/4 |
| | $6\times10^{-10}$ | 0/4 |
| | $10^{-10}$ | 0/4 |
| | $3\times10^{-11}$ | 0/4 |
| sEVs | $10^{-5}$ | 4/4 |
| | $2\times10^{-6}$ | 4/4 |
| | $4\times10^{-7}$ | 3/4 |
| | $8\times10^{-8}$ | 0/4 |
| | $2\times10^{-8}$ | 1/4 |
| | $3\times10^{-9}$ | 0/4 |
| | $6\times10^{-10}$ | 0/4 |
| | $10^{-10}$ | 0/4 |
| | $3\times10^{-11}$ | 0/4 |
| cell lysate (negative) | $10^{-5}$ | 0/4 |

PK1-N2a-RML 1st isolation

| Sample | Dilution | Positive wells |
|---|---|---|
| cell lysate | $10^{-5}$ | 4/4 |
| | $2\times10^{-6}$ | 4/4 |
| lEVs | $10^{-5}$ | 4/4 |
| | $2\times10^{-6}$ | 4/4 |
| | $4\times10^{-7}$ | 4/4 |
| | $8\times10^{-8}$ | 4/4 |
| | $2\times10^{-8}$ | 4/4 |
| | $3\times10^{-9}$ | 2/4 |
| | $6\times10^{-10}$ | 0/4 |
| | $10^{-10}$ | 1/4 |
| | $3\times10^{-11}$ | 0/4 |
| sEVs | $10^{-5}$ | 4/4 |
| | $2\times10^{-6}$ | 4/4 |
| | $4\times10^{-7}$ | 4/4 |
| | $8\times10^{-8}$ | 3/4 |
| | $2\times10^{-8}$ | 4/4 |
| | $3\times10^{-9}$ | 2/4 |
| | $6\times10^{-10}$ | 1/4 |
| | $10^{-10}$ | 0/4 |
| | $3\times10^{-11}$ | 0/4 |
| cell lysate (negative) | $10^{-5}$ | 0/4 |

PK1-N2a-RML 2nd isolation

| Sample | Dilution | Positive wells |
|---|---|---|
| cell lysate | $10^{-5}$ | 4/4 |
| | $2\times10^{-6}$ | 4/4 |
| lEVs | $10^{-5}$ | 4/4 |
| | $2\times10^{-6}$ | 4/4 |
| | $4\times10^{-7}$ | 4/4 |
| | $8\times10^{-8}$ | 4/4 |
| | $2\times10^{-8}$ | 4/4 |
| | $3\times10^{-9}$ | 2/4 |
| | $6\times10^{-10}$ | 0/4 |
| | $10^{-10}$ | 0/4 |
| | $3\times10^{-11}$ | 1/4 |
| sEVs | $10^{-5}$ | 4/4 |
| | $2\times10^{-6}$ | 4/4 |
| | $4\times10^{-7}$ | 4/4 |
| | $8\times10^{-8}$ | 2/4 |
| | $2\times10^{-8}$ | 1/4 |
| | $3\times10^{-9}$ | 1/4 |
| | $6\times10^{-10}$ | 0/4 |
| | $10^{-10}$ | 1/4 |
| | $3\times10^{-11}$ | 0/4 |
| cell lysate (negative) | $10^{-5}$ | 0/4 |

**Supplementary Table S2**: Summary of detection of prion converting activity in cell lysates from CAD5-RML or N2a-PK1-RML cells (positive control) and isolated lEVs and sEVs from these cells. The negative control was cell lysate from CAD5 or N2a-PK1 cells. Each sample was serially diluted in quadruplicate. The lowest dilution where at least two wells from quadruplicates (dark green) marks the SD50.
